# The protein structurome of *Orthornavirae* and its dark matter

**DOI:** 10.1101/2024.09.25.615016

**Authors:** Pascal Mutz, Antonio Pedro Camargo, Harutyun Sahakyan, Uri Neri, Anamarija Butkovic, Yuri I. Wolf, Mart Krupovic, Valerian V. Dolja, Eugene V. Koonin

**Author notes:** Correspondence to: Pascal Mutz, Valerian V. Dolja or Eugene V. Koonin.

## Abstract

Metatranscriptomics is uncovering more and more diverse families of viruses with RNA genomes comprising the viral kingdom *Orthornavirae* in the realm *Riboviria*. Thorough protein annotation and comparison are essential to get insights into the functions of viral proteins and virus evolution. In addition to sequence- and hmm profile-based methods, protein structure comparison adds a powerful tool to uncover protein functions and relationships. We constructed an *Orthornavirae* ‘structurome’ consisting of already annotated as well as unannotated (‘dark matter’) proteins and domains encoded in viral genomes. We used protein structure modeling and similarity searches to illuminate the remaining dark matter in hundreds of thousands of orthornavirus genomes. The vast majority of the dark matter domains showed either ‘generic’ folds, such as single α-helices, or no high confidence structure predictions. Nevertheless, a variety of lineage-specific globular domains that were new either to orthornaviruses in general or to particular virus families were identified within the proteomic dark matter of orthornaviruses, including several predicted nucleic acid-binding domains and nucleases. In addition, we identified a case of exaptation of a cellular nucleoside monophosphate kinase as an RNA-binding protein in several virus families. Notwithstanding the continuing discovery of numerous orthornaviruses, it appears that all the protein domains conserved in large groups of viruses have already been identified. The rest of the viral proteome seems to be dominated by poorly structured domains including intrinsically disordered ones that likely mediate specific virus-host interactions.

**IMPORTANCE:** Advanced methods for protein structure prediction, such as AlphaFold2, greatly expand our capability to identify protein domains and infer their likely functions and evolutionary relationships. This is particularly pertinent for proteins encoded by viruses that are known evolve rapidly and as a result often cannot be adequately characterized by analysis of the protein sequences. We performed an exhaustive structure prediction and comparative analysis for uncharacterized proteins and domains (‘dark matter’) encoded by viruses with RNA genomes. The results show the dark matter of RNA virus proteome consists mostly of disordered and all α-helical domains that cannot be readily assigned a specific function and that likely mediate various interactions between viral proteins and between viral and host proteins. The great majority of globular proteins and domains of RNA viruses are already known although we identified several unexpected domains represented in individual viral families.

## Introduction

Viruses are the most abundant biological entities on earth infecting all life forms. In the recently adopted comprehensive taxonomy, all viruses have been divided into 6 realms one of which, *Riboviria*, consists of an enormous variety of viruses with RNA genomes that encode homologous replication enzymes, RNA-dependent RNA polymerase (RdRP), in the kingdom *Orthornavirae*, or reverse transcriptase, in the kingdom *Pararnavirae*. In the last few years, metatranscriptomics along with targeted approaches have been uncovering diverse orthornavirus families at an increasing pace (1-4). Orthornaviruses have small genomes, mostly, between 3-20 kilobases (kb), with the upper bound of 35-64 kb in nidoviruses (5, 6), and accordingly, encode limited repertoires of protein domains. Thorough annotation and comparison of viral proteins provides ample insights into protein functions, virus evolution and in some cases, virus-host association. The RdRP is the only protein that is conserved in all orthornaviruses (7) and thus serves as the primary query for the discovery of orthornaviruses in metatranscriptomes (1, 8, 9). Other proteins conserved in large groups of orthornaviruses include helicases and proteases of different families, mRNA capping enzymes, and capsid proteins of different types. Several other protein domains are conserved across more narrow ranges of orthornaviruses, often associated with specific host organisms. Such domains include the movement proteins (10) and AlkB family oxygenases (11) found in a variety of plant viruses, ADP-ribose-binding Macro domains encoded by several families of animal viruses, lysozymes encoded by a variety of RNA bacteriophages (1, 12), and more. All these protein domains are conserved at the sequence level so that annotation using protein family profiles can delineate their core regions in viral genomes with high confidence. However, even exhaustive (viral) protein profile generation and searches leave many unannotated proteins and large protein regions in newly discovered orthornaviruses. The question thus remains how these unannotated portions of viral proteins are related to each other and what are their structures and potential functions.

Protein structures are in general far more strongly conserved than sequences, and therefore, structural comparisons have the potential to illuminate the ‘dark matter’ of viral proteomes as compellingly demonstrated for proteomes of cellular life forms, in particular, the human proteome (13, 14). With the advent of high-accuracy protein structure prediction methods, such as AI-based AlphaFold2 and RosettaFold or using transformer protein language models such as ESMfold, comprehensive protein structure prediction and analysis have become realistic (15-17). Large-scale databases of protein structure models are already available (e.g. EBI: https://alphafold.ebi.ac.uk/ (18), ESM Atlas: https://esmatlas.com/ESM atlas (17)), covering either single species proteomes or metatranscriptomes such as ESM atlas. However, no such databases were available for viral proteins while conducting this research, in part, due to the difficulty of processing polyproteins that are encoded by many viruses, in particular, members of *Orthornavirae*. Only recently, Nomburg and colleagues presented predicted structures for proteins of eukaryotic viruses (19).

We have previously demonstrated the utility of protein structure prediction using AF2, followed by comparison to structure databases, for predicting functions of uncharacterized proteins or protein domains of DNA viruses, revealing, in particular, multiple cases of exaptation of host enzymes (20, 21). We were then motivated to explore in depth the proteomic ‘dark matter’ of orthornaviruses using a similar approach. To this end, we used a previously published dataset of (predicted) orthornavirus genomes spanning over 300,000 viral contigs from nearly 500 (operational) virus families discovered in metatranscriptomics (1). Of note, nearly 400 virus families are operational and not formally ratified. We pre-processed the predicted viral proteins to isolate unannotated domains and hypothetical open reading frames (ORFs), modeled them in addition to well annotated domains using AF2 and ESMfold, and performed structure comparisons among all viral proteins to compose a virus protein ‘structurome’. These structural models were then compared to structures of cellular proteins represented in the PDB. We found that the vast majority of unannotated regions of orthornaviral proteins and smaller unannotated ORFs were either predicted to form a ‘generic’ fold, such as a single α-helix, or could not be modeled with high confidence, suggesting non-globular structure. Nevertheless, several globular domains, mostly, represented in one or more narrow viral lineages and not previously detected either in any orthornaviruses or at least in a given viral family were predicted. Taken together, the results of this analysis indicate that the widespread globular domains comprising the proteome of orthornaviruses are largely known whereas newly identified proteins and domains are lineage-specific, are in many cases non-globular, and are likely to be involved in interactions between viral and host proteins.

## Results

### Annotated and unannotated domains and proteins of orthornaviruses

In order to compile a comprehensive set of orthornavirus proteins, we used a recently published dataset (hereafter environmental metatranscriptome RNA virus (EMRV) set) of predicted orthornavirus genomes spanning more than 370,000 viral contigs from nearly 500 operationally defined virus families of which 98 had been approved by the ICTV at the time of publication (2022) (1). All proteins and domains annotated in this study were extracted, yielding a set of 647,383 protein sequences. Then, evolutionarily conserved but unannotated (putative) proteins and domains that are conserved in groups of viruses were identified (see Suppl.Fig.1 for a schematic and Methods for details). In brief, ORFs from start to stop were predicted in all 6 frames and matched to the published annotations (1). ORFs without annotation and with only partial annotation were processed to retrieve the unannotated sequence stretches. In the case of polyproteins and multidomain proteins, unannotated and partially annotated proteins of at least 200 amino acids (aa), in which a continuous stretch of at least 60 aa was unannotated, hereinafter conserved unannotated domains (CUDs), were extracted. Unannotated ORFs between 60 and 199 aa (n=1,362,871) were not sliced further and kept for downstream analysis (hereinafter ‘ORFans’). To identify evolutionarily conserved sequences that are likely to be expressed and functional, ORFans and CUDs were clustered by sequence similarity (see Methods). All clusters that included proteins from genomes covered by at least three leaves in the RdRP-based phylogenetic trees or at least one CUD were retained for further analysis, resulting in 6,117 clusters of ORFans representing 100,694 sequences and 13,085 CUDs representing 31,247 sequences (Suppl. Fig.2 A, B).

**Fig. 1:**
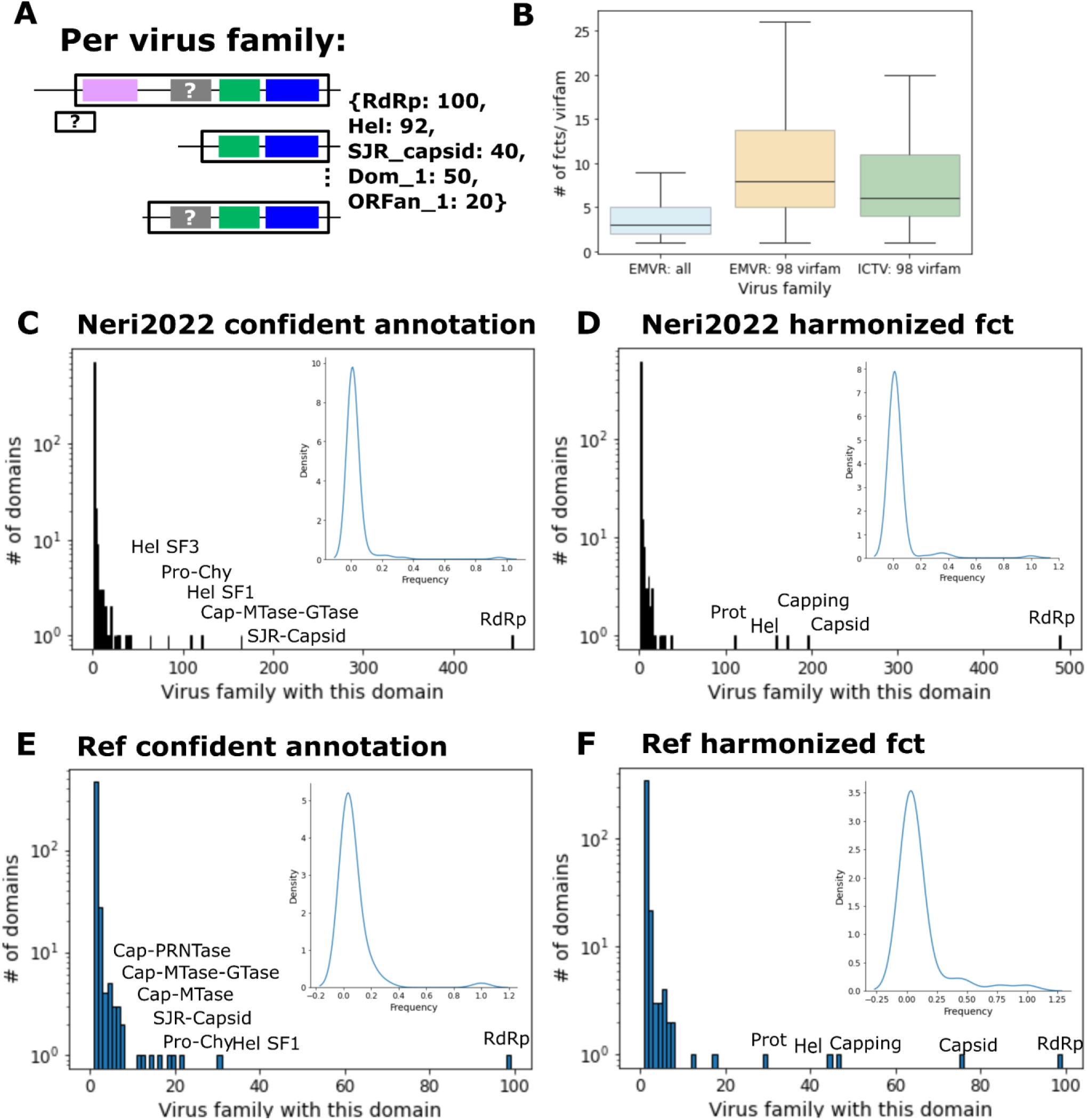
Frequencies of EMRV profile annotated vs ICTV exemplar annotated proteins across virus families. **(A)** Schematic of a prototype pangenome per virus family with 100 contigs which are either incomplete or complete but all contain the RdRp, and some also contain Helicase (Hel), a single jelly-roll capsid (SJR_capsid), an un-annotated domain (Dom_1, CUD) or an ORFan (ORFan_1). **(B)** Number of unique annotated domains per virus family for all virus families in the EMRV set (‘EMRV: all’), of the 98 named virus families with a corresponding family in the ICTV exemplar set (‘EMRV: 98 virfam’) and of the ICTV exemplar set (‘ICTV: 98 virfam’). **(C)** Number of unique annotations based on profile comparison per virus family across all 498 families. **(D)** Same as C but with harmonized functions (e.g. combining all Helicase related labels as ‘Hel’). **(E)** Number of unique annotations within the ICTV exemplar virus families based on nvpc profile db comparison. **(F)** Harmonized functions (e.g. ‘capsid’ represents the functional tags ‘nucleoprotein’, ‘SJR capsid’, ‘core’ and others assigned to capsid and nucleocapsid proteins) across the ICTV exemplar virus families as in E together with proteins which are annotated in GenBank but not in nvpc. **(C-F)**: Inset shows frequencies for all functional domains that are present in at least 2 families.

**Fig. 2:**
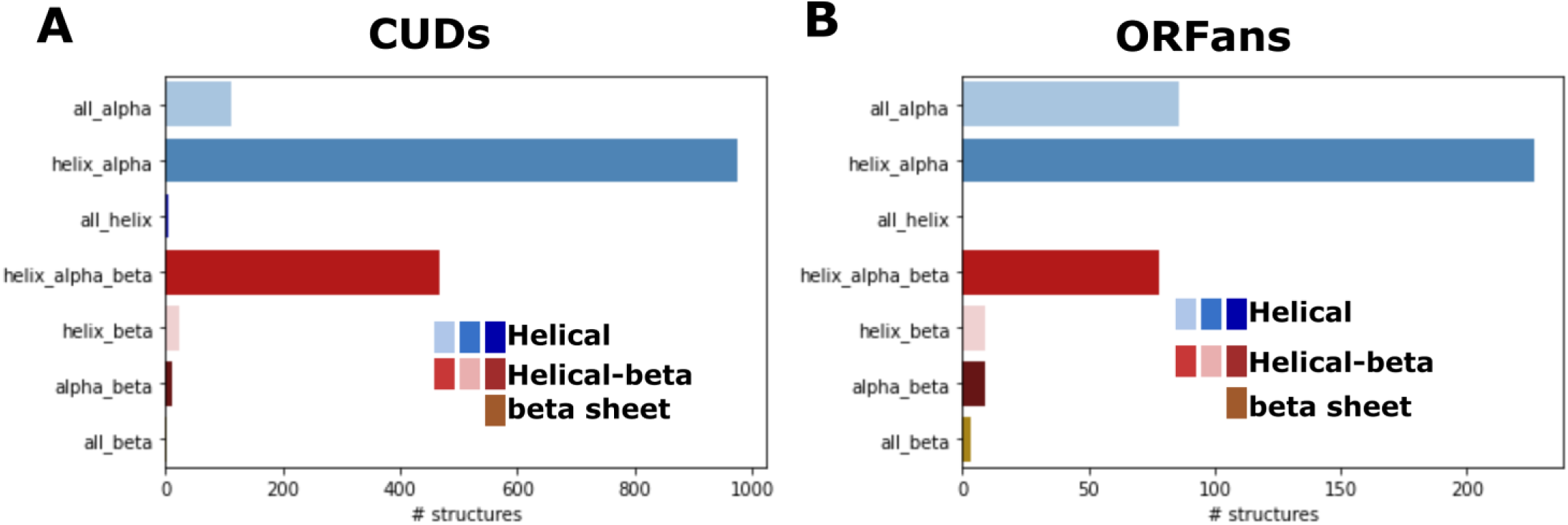
Secondary structure assignments for CUDs and ORFans. Psique-based secondary structure assignment are shown for all CUDs **(A)** and ORFans **(B)** with a mean plddt >= 70. Helical types in blue color range, beta-strand and helical in red color range, all-beta in brown and other in grey.

Next, we assessed which known protein domains were missed during profile-based annotation and might be present within the CUD and ORFan set. To this end, we downloaded a curated set of viral reference genomes from the ICTV (ICTV exemplars, https://ictv.global/vmr, VMR_MSL38_v2) matching the 98 virus families recognized in the EMRV set and extracted the annotated domains and proteins (n=32,648) from the GenBank files (hereafter exemplar domains). Exemplar domains were clustered and compared against the same viral profile databases used for EMRV annotation ran using hhsearch (22). Clusters with at least one highly confident hit (probability ≥ 95%) were harmonized and assigned accordingly. As a result, 64% of ICTV clusters were assigned confidently, spanning 83% of all exemplar domains. Across virus families, about 70% of all clusters associated with a given family could be confidently assigned, increasing to 80% if considering only clusters spanning domains on the RdRP-encoding segment, those recovered by Neri *et al.* (1) and included in the EMRV data set (Suppl. Fig. 3 A). The clusters without confident assignment were dominated by functionally uncharacterized domains (Suppl. Fig. 3 B). Thus, some known viral proteins and domains are not identified with high probability by the used viral protein profiles and might be present among the CUDs and ORFans. These under-annotated domains were identified as part of the refinement of orthornavirus proteome as described below.

### The pan-proteome of *Orthornavirae*

Combining the annotated domains and proteins with CUDs and ORFans, we constructed a pan-proteome for each orthornavirus family. All domains and ORFs (annotated or not) were clustered by sequence similarity and domains were either labeled by their functional tag or as CUD or ORFan. Similar procedure was performed for the ICTV exemplar set, and the resulting virus family pan-proteomes were compared (Fig.1 A). For the 98 virus families represented in both the ICTV exemplars and the EMRV set, comparable numbers of functions (profile assignments) per virus family were detected, confirming that robust domain annotation was obtained for the EMRV set. Functional assignment per virus family across all 498 families was lower because about 25% of the families included viruses for which only the RdRP was detected, that is, most likely, viruses with segmented genomes (Fig.1 B).

A substantial majority of the domains annotated by a particular profile were found in a single virus family in both the EMRV set (85%) and the ICTV exemplar set (89%). Thus, these domains represent virus family- or even genus- or species-specific functions (Fig. 1 C,E). The RdRP was the only protein represented in all virus families. Very few other domains and functions were found to be broadly distributed across virus families. These widespread functions are different types of capsids (40% of families in the EMRV set vs 76% in the ICTV exemplar set), mRNA capping enzymes (35% vs 47%), helicases (33% vs 45%) and proteases (22% vs 30%) (Fig. 1 D,F). The consistent lower abundance of these domains across virus families within the EMRV set compared to the ICTV exemplars is probably due to the absence of non-RdRP-encoding segments of multipartite genomes in the EMRV set.

### The structurome of *Orthornavirae*

To predict structures and functions of the CUDs and ORFans, we generated structural models using AlphaFold2 (15). Only a fraction of structures (about 34% of CUDs and 10% of ORFans) could be predicted with high confidence (mean plddt score ≥70), indicating the possibility of larger intrinsically disordered stretches (Suppl. Figs. 4 A and 5 A). To address the first possibility, we predicted intrinsically disordered structures in CUDs and ORFans. Although we found no significant correlation with the plddt score of the structural models, proteins predicted to have disordered regions have mostly a plddt score below 70 (Suppl. Figs. 4 B and 5 B). To account for the uncertainty of correct folds among low plddt structures, we clustered all CUDs and ORFans independently with FoldSeek (0.8 coverage) and kept only clusters in which at least one member had a mean plddt score of 70 or higher, resulting in 412 ORFans and 1,594 CUDs.

These representatives were compared to PDB using Dali (23). Secondary structure fold prediction with Psique (24) indicated an all-α-helical fold for about 69% of CUDs and nearly 76% of ORFans (Fig. 2). Further, for 53% of the CUDs and 35% of the ORFans, folds distinct from the simplest ones, such as a single α-helix or helix-turn-helix (HTH), were predicted. Those were considered as CUD and ORFan of interest (COI and OOI hereafter, respectively). About 37 % and 13 % of the ORFans and CUDs with a predicted simple fold were predicted to contain at least one transmembrane domain compared to about 8 % and 9 % of the more structured ORFans and CUD, respectively, indicating a substantial enrichment of small transmembrane proteins among ORFans with a simple fold. Most of the COIs and OOIs were confined to a single virus family (Supp. Fig. 6). FoldSeek clusters spanning representatives of multiple families represent mainly capsid proteins (see below). Further, COIs were checked with a ‘neighborhood’ approach to determine whether a given COI was confidently annotated in a related genome from the same virus family, pointing to its function (Fig. 3, A,B). In brief, COIs were searched against the annotated domains using psi-blast to obtain a provisional annotation of the COI. Next, related proteins from neighboring genomes with or without COI were aligned and the respective annotations were mapped to the alignment. Whenever annotations from neighboring proteins overlapped with the COI, the annotation was compared with the psi-blast hit. If consistent, the COI was considered a ‘refinement’ and was not investigated further. Whenever there was no overlap with an annotated domain, a conflicting or mixed result or no psi-blast hit, the COI was kept for manual inspection of the Dali results. Of the 852 COIs with confidently predicted structures, 62 were from genomes not phylogenetically assigned, 553 represented ‘refinements’, many as part of the RdRP.

**Fig. 3:**
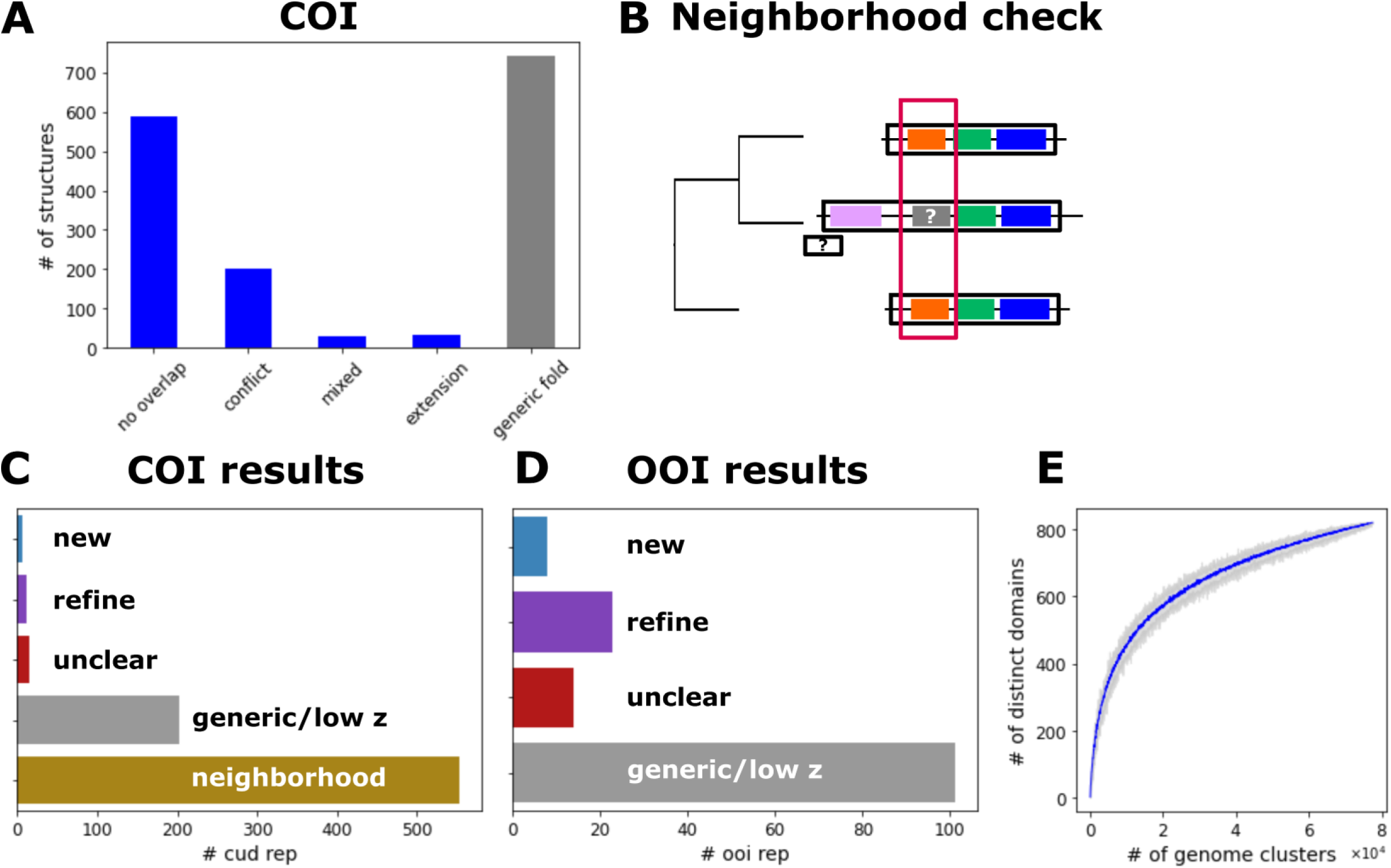
Overview of domains and ORFans of interest. **(A)** Number of representative COI (conserved unannotated domain of interest) structures binned as follows: i) no overlap with present annotation in genome; ii) conflict: there is a present annotation that slightly overlaps with the provisional CUD annotation; iii) mixed: members of a CUD cluster had substantially different provisional PSI-BLAST annotations; iv) extension: the provisional psi-blast annotation of a CUD extended the annotation of an existing profile-based annotated domain; v) generic fold: based on Dali results, the fold is a single helix, HTH, a beta-hairpin or disordered. Categories i-iii were analyzed further. **(B)** Schematic of the neighborhood analysis. Homologous multidomain proteins or polyproteins of neighboring genomes were aligned, and protein annotations were mapped on the alignment. If a putative COI region overlapped a confident annotation, it was considered annotated. **(C)** Number of COIs which were considered annotated as a result of the neighborhood analysis (bottom bar) and results of the semi-manual examination of COIs. New: a COI representative with a predicted structure not reported previously for the given virus family. Refine: Dali results pointed to a refinement of the annotation as the structure/function was already reported for other members in this virus family. Unknown: a high confidence model was obtained for a COI but Dali hits were inconclusive (mainly, alpha/beta domains). Generic/ low z: the structure is too generic to produce meaningful Dali hits (e.g. an alpha helix with a small beta hairpin) or the Dali z-score was no significant (below 4). **(D)** Results of the semi-manual check of OOIs. Binning is as in C. **(E)** Rarefaction curve of distinct domains as function of the number of sampled genome clusters (leaves). Blue line represents mean of 30 bootstraps and grey area shows the range of unique domains at each sampling step (step size: 50 genome clusters).

To incorporate the predicted structures of COIs and OOI into the context of known structures across all *Orthornavirae* families, we constructed a structurome of *Orthornavirae* by modeling one representative structure per functional tag per virus family from the pangenome (e.g. one representative RdRP structure per virus family, 2,421 annotated representatives in total). The ICTV exemplar domains lacking strong profile annotation were modeled too (3,000 representatives). This modeling resulted in 6,419 predicted structures which were pre-clustered with FoldSeek (0.8 coverage, 4,022 clusters). The great majority of the structures, 88%, remained singletons (Suppl. Fig. 7 A). In 27 cases, OOIs were found in a FoldSeek cluster together with annotated domains and/or ICTV exemplar domains, and the same was found for 13 COIs. Then, 4,022 cluster representatives were analyzed by an all-vs-all Dali comparison, finalizing the *Orthornavirae* structurome. Structures within the structurome were clustered based on the Dali all-vs-all z-scores in an iterative procedure in which individual structures were added to a cluster as long as the mean z-score to each other structure already present was above a given z score. The threshold was set at z-score of 7 or higher (‘z7 clusters’) to avoid over-clustering. This procedure resulted in 333 z7 clusters of which 59% contained 2 structures whereas the largest cluster consisted of 43 structures (Suppl. Fig. 7B). These 333 z7 clusters represented 1201 (30%) of all representative structures in the Dali all-vs-all run. The structurome was visualized as structure-similarity network in which each structure is a node connected to other nodes via edges weighted by the pairwise z-score. Suppl. Fig. 8 shows a subset of the network in which only structures present in a z7 cluster are shown or which have at least one connection to a structure within a z7 cluster with a z-score of 7 or higher (about 36% of all structures in the Dali all-vs-all run). As expected, we detected z7 clusters of structures coming from well known, functionally characterized domains such as RdRP, helicases, single jelly-roll (SJR) capsid proteins and proteases. Some functional labels were distributed across several z7 clusters. For example, RdRP structures which dominate the structurome (489 structures initially) contributed to 4 z7 clusters with one harboring about 88 % of the representative RdRP structures. (Suppl. Fig. 7B).

About 28% (n=92) z7 clusters consisted of OOIs and COIs only, that is, represented unique shared folds. Notably, nearly all α-helical domains and proteins (annotated or not) were placed in eight larger z7 clusters (10 representatives or more) in which they are connected by z-scores of 7 or higher but these connections apparently reflect generic structural similarity rather than shared distinct folds (hubs labeled ‘Helical’ in Suppl. Fig. 8).

OOIs and COIs were inspected semi-manually by considering Dali results, secondary structures and structural relationships within the structurome. All-helical COIs and OOIs mostly did not show conclusive Dali hits, with moderate structural similarity between small α-helical modules detected across apparently unrelated proteins. Inspection of COIs and OOIs predicted to fold into globular structures either revealed a fold and function not yet reported for a given virus family (‘new’), or a fold not yet reported for a given virus family for which no clear function could be assigned due to significant but too variable Dali hits (‘unclear’), or indicated a refinement of the current profile based annotation of a given virus family (‘refined’). Altogether, we classified the predicted structures for OOIs and COIs, respectively, as follows: 8 and 7 new; 14 and 16 unclear; 23 and 12 refined (Fig. 3 C, D).

Saturation of the rarefaction curve of distinct domains sampled randomly across *Orthornavirae* genome clusters suggests that the substantial majority of widely distributed *Orthornavirae* domains are already known (Fig. 3 E). Conversely, new viral domains are expected to be found in single virus families which is in line with the findings of this study. Having both the structure-based and profile-based annotations at hand, we aimed to identify the core set of unique domains per virus family shared by at least 50% of genomes (see Methods for details) and compare it to the overall distribution of domains per virus family. For most orthornavirus families, we identified a small core, often consisting of the RdRp alone, and a ‘shell’ of domains with intermediate frequencies (Suppl. Fig. 19). This distribution of domain frequencies could reflect true variation across a virus family, e.g. on the genus level, but also the presence of incomplete genomes in the dataset, in particular, those with multiple segments, and a lack of complete domain annotation.

### Discovery of new domains in the orthornaviral structurome

‘Unexpected’ OOIs and COIs identified in a particular virus family potentially could be either truly novel, that is, not reported so far in any *Orthornavirae*, or new to the given virus family. There were no unequivocal examples of the former case although we identified two OOIs from the putative family *f.0145* (o.0036, c.0025, *Kitrinoviricota*) with a predicted β-barrel fold. Best but not consistent Dali hits were to bacterial β-barrel fold proteins (top z-scores ∼5, PilZ domain and HCP3, a paralogue of type VI secretion system effector, Hcp1, pfam PF05638, for the two OOIs, respectively) and no structural similarity was observed to known virus proteins in the *Orthornavirae* structurome (Suppl. Fig. 9).

Apart from these two β-barrel domains for which there were no homologs in *Orthornavirae* (or, to our knowledge, any other viruses), we identified several other domains known to be encoded by *Orthornavirae* members but here found in unexpected virus families. Several COI and OOI products with different folds seem to be involved in nucleic acid binding. One of these nucleic acid-binding domains is the phytoreovirus core-P7 dsRNA-binding domain (P7-dsRBD) that is known to be encoded by members of *Sedoreoviridae (25)*and here was found in proteins of various virus families as annotated by profile comparison: *Chrysoviridae* (70/81 leaves covered), *Endornaviridae* (30/222), *Megabirnaviridae* (3/12), *f.0281.base-Megabirna* (7/29), *f.0285* (31/94), *Flaviviridae* (4/360), with z-scores of 8-11 to each other (see pangenome (Suppl. Table 1) for all virus families with this domain). We identified an OOI with a similar fold in *Picobirnaviridae* (in genomes of 14/1376 leaves scattered across the tree; Fig.4). Structure comparison against PDB revealed prominent similarity to nucleotide monophosphate (NMP) kinases, such as adenylate kinase (z-score ∼8). Thus, proteins with a core-P7-like RBD fold likely originated from cellular NMP kinases and might have spread horizontally among diverse viruses. Representative viral protein structures with P7-dsRBD from different virus families including *Phytoreovirus*, *Chrysoviridae* and *Picobirnaviridae* were aligned with homologous cellular kinases using FoldMason (26) and a phylogenetic tree was constructed using IQTree2 (27). Most of the viral P7-dsRBDs formed a distinct clade separate from the cellular kinases (Fig. 4C) which is compatible with functional divergence after exaptation and subsequent horizontal spread. Exceptions are the P7-dsRBDs of *Picobirnaviridae* which clustered within the cellular kinase clades, suggestive of independent exaptation events (Fig. 4C). While the Walker A motif is intact in viral core-P7-like RBD fold proteins, the Walker B motif is missing, indicating loss of kinase activity (Suppl. Fig. 10). The P7-dsRBD domains are found either as stand-alone proteins or are incorporated into viral polyproteins (Suppl. Fig. 12). This domain was identified in 3 families from the order *Ghabrivirales* (*Chrysoviridae*, *f.0296*, *f.0285*) indicative of an old acquisition but in contrast seems to have been more recently acquired at least twice by *Picobirnaviridae* members (Fig. 4B, C). Other members of *Picobirnaviridae*, predicted to infect bacterial hosts, have been shown to encode a putative lysozyme (completely unrelated to dsRBD or kinases) in the same genomic location (1). Furthermore, yet other members of *Picobirnaviridae* encode a capsid protein 5’ of the RdRP ORF, demonstrating the flexibility of *Picobirnaviridae* to capture diverse ORFs 5’ of the RdRP ORF. The exact functions of the P7-dsRBD domains in different virus families remain unclear. Notably, all experimentally characterized viruses from families with P7-dsRBD domains have dsRNA genomes, suggesting that this domain is specifically involved in capsid-associated dsRNA transactions (given that all viruses produce replicative dsRNA intermediates in the host cell). It seems likely that these domains are involved in transcription and/or RNA packaging as reported for the phytoreovirus P7 (28), but additional or alternative roles, for example, in the suppression of the host RNAi system, cannot be ruled out.

**Fig. 4:**
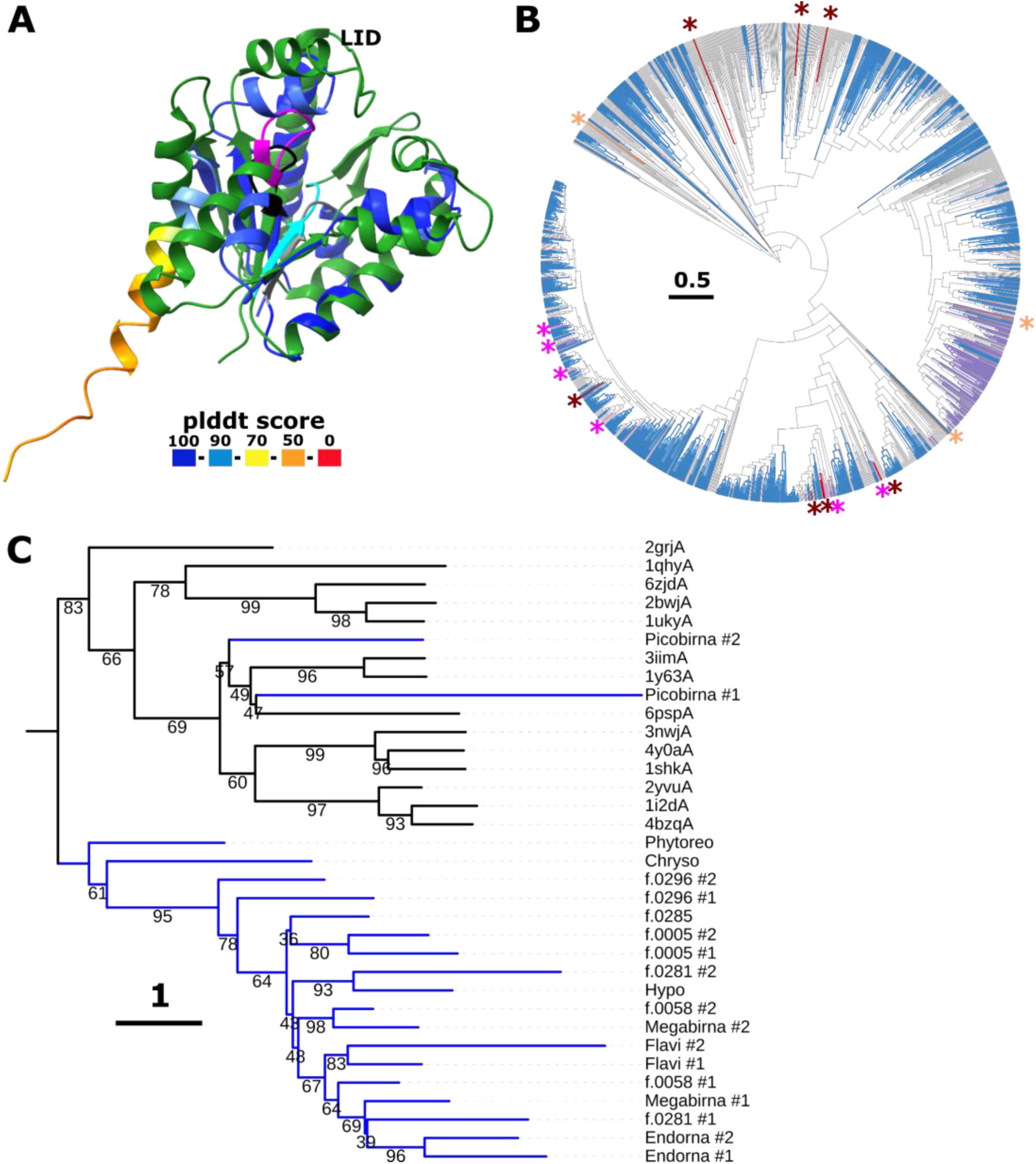
Phytoreovirus core-P7 dsRNA-binding domain with a kinase fold in orthornaviruses. **(A)** Superposition of *Picobirnaviridae* 5’ ORFan (colored by plddt score, Walker A motif is shown in black, position of degraded Walker B motif is shown in grey) with adenylate kinase from *Methanococcus igneus* (6psp, green, Walker A motif shown in magenta, Walker B motif is shown in cyan, z-score 9.1). **(B)** Phylogenetic distribution of contigs encoding P7-dsRBD (red branches and asterisks), lysozyme (orange and asterisk) and capsid protein (purple) 5’ of the RdRp within *Picobrinaviridae*. Blue color indicates contigs containing less than 180 nt in front of the RdRp ORF (likely incomplete). **(C)** Phylogenetic tree based on a structure-guided alignment of viral P7-dsRBD domains found in different virus families by structure comparison (Picobrinaviridae) or profile comparison (EMRV set) with structurally similar kinases (z-scores 6-11; order as in tree: Dephospho-CoA kinase from *Thermotoga maritim*a (2grjA); Chloramphenicol phosphotransferase from *<u>Streptomyces venezuelae</u>* (1qhyA); Adenylate kinase 3 from *Homo sapiens* (6zjdA); Adenylate kinase 5 from *Homo sapiens* (2bwjA); Uridylate kinase from Saccharomyces cerevisiae (1ukyA); Atypical mammalian nuclear adenylate kinase hCINAP from Homo sapiens (3iimA); Probable kinase from *Leishmania major Friedlin* (1y63A); Adenylate kinase from *Methanococcus igneus* (6pspA); Shikimate kinase from *Arabidopsis thaliana* (3nwjA); Shikimate kinase from *Acinetobacter baumannii* (4y0aA); Shikimate kinase from *Erwinia chrysanthemi* (1shkA); APE1195 from *Aeropyrum pernix* K1 (2yvuA); ATP Sulfurylase from *Penicillium chrysogenum* (1i2dA); APS kinase CysC from *Mycobacterium tuberculosis* (4bzqA);). Branches of viral P7-dsRBD are colored in blue, and those of cellular kinases are colored in black.

An OOI containing a common RBD fold (29) consisting of four β-strands and two α-helices (obviously, unrelated to the kinase-derived RBD discussed above) was identified in members of the *Hepeviridae* family (Fig. 5). This OOI is located 3’ of the non-structural polyprotein and capsid protein ORFs. It is unclear whether this OOI actually binds RNA because the top Dali hits are the RBD that have lost the RNA-binding capacity and are instead involved in protein-protein interactions, such as RBD2 of the *Arabidopsis* protein HYL1 (30).

**Fig. 5:**
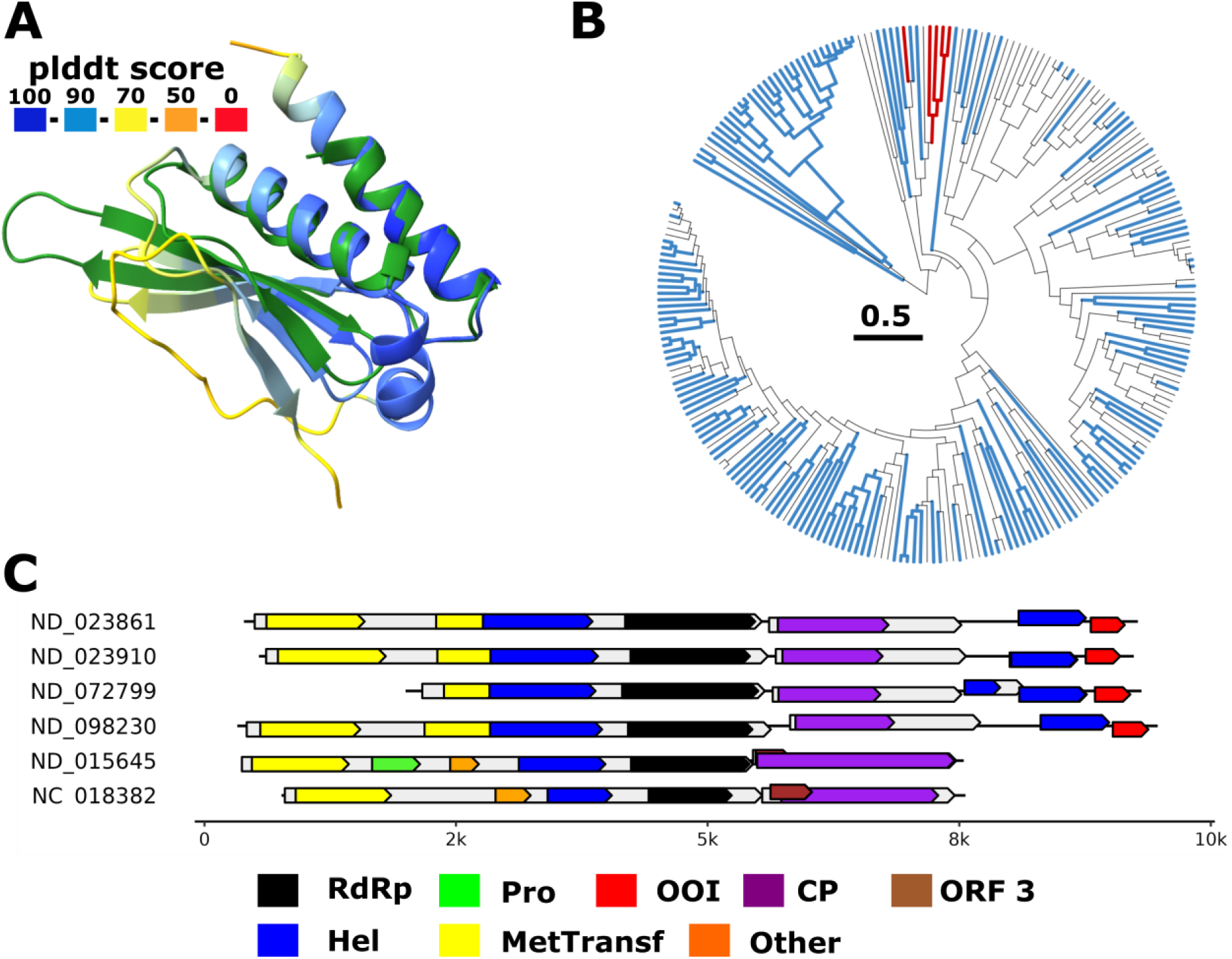
Inactive RNA-binding domain fold in *Hepeviridae*. **(A)** Superposition of likely inactive RNA binding domain (RBD)-fold found in *Hepeviridae* (colored by plddt score) with RBD 2 from *A. thaliana* protein HYL1 (pdb 3adj, green, z-score 7.5). **(B)** Phylogenetic tree of *Hepeviridae* RdRp. Branches containing the OOI with RBD-fold are colored in red. Branches with no coding capacity after capsid are colored in blue. **(C)** Genome maps for representative *Hepeviridae* members encoding (or not) for the OOI. Annotations are based on profile analysis (1) and GenBank annotation (NC 018382). Protein domains: RdRp, RNA-dependent RNA polymerase, Hel, Helicase, CP, capsid protein, Pro, Protease, ORF3, *Hepeviridae* ORF 3, Other: additional *Hepeviridae* domains, OOI: ORFan of interest.

Another nucleic acid binding domain is a dsRBD that is found in many virus families such as *Nodaviridae*, *Astroviridae* and *Reoviridae* and functions as viral suppressor of host RNAi defense (see virus family pangenomes). We additionally identified related dsRBD as a COI domain in *f.0092*, basal to *Permutotetraviridae* (Suppl. Fig. 12). This domain was detected in genomes represented by nearly all leaves of this family (16/17).

Yet another fold implicated in nucleic acid binding is a winged helix-turn-helix (wHTH) domain found in family *f.0008* basal of *Polycipiviridae* (Fig.6). The wHTH domain apparently was acquired recently by *f.0008* members as it was only found in a distinct, distal clade (Fig. 6 C). In addition to the wHTH domain, the same and additional *f.0008* family members were shown to encode an SJR-fold protein (Fig. 6 B-D). Given the genome architecture of *f.0008* members with an annotated larger capsid ORF 5’ of the newly identified SJR protein, and given that it clusters with other plant movement proteins (MP) in the structurome, it is highly likely that this is a 30K superfamily MP that evolved by duplication of a SJR capsid protein followed by neofunctionalization (10). Thus, plants are likely the hosts for at least this clade of *f.0008* members.

**Fig. 6:**
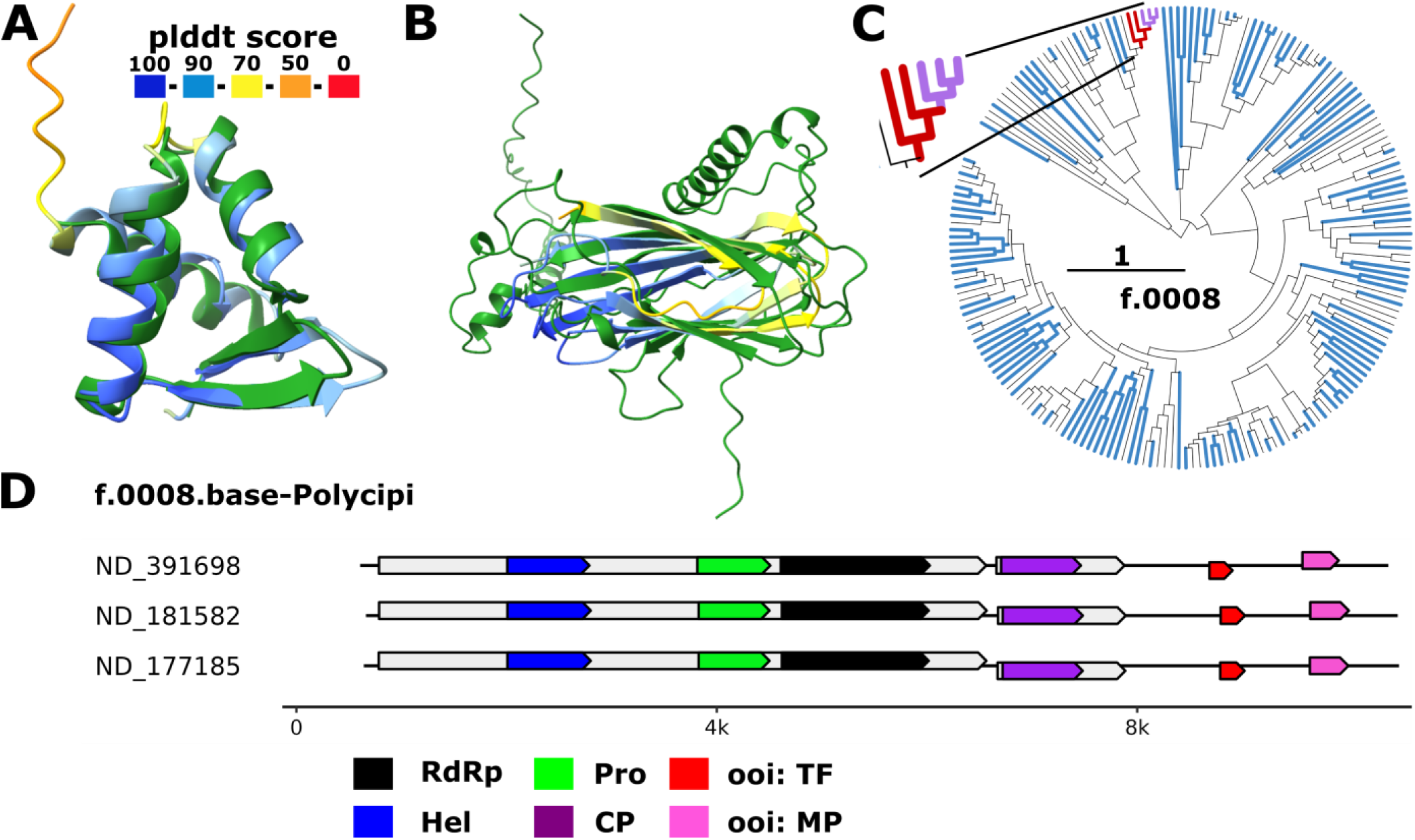
Winged helix-turn-helix domain and movement protein ORFans in a novel viral family. **(A)** Superposition of wHTH domain identified in *f.0008.base-Polycipi* (plddt score colored) with mouse HOP2 DNA-binding wHTH domain (2mh2, green, z-score: 11.1). **(B)** Superposition of predicted movement protein encoded by *f.0008* members (plddt colored) with an annotated movement protein domain from *Betaflexiviridae* (AlphaFold2 modeled, green, MP_30K, z-score: 7). **(C)** Phylogenetic distribution of contigs encoding only the predicted movement protein (red) or both movement protein and wHTH (purple). Blue branches indicate contigs with less than 180 nt after the capsid encoding ORF which are likely incomplete. **(D)** Representative genome maps for members of *f.0008* carrying the respective ORFs of interest (OOI). Protein domains: RdRp, RNA-dependent RNA polymerase, Hel, Helicase, CP, capsid protein, Pro, Protease, OOI, ORF of interest (wHTH or MP).

Independently of the putative MP and wHTH, other members of the family *f.0008* were found to encode a galactose-binding domain (Suppl. Fig. 13, Dali z-score of 15.5 against a bacterial carbohydrate binding domain). The general genome organization of *f.0008* members differs between those encoding the wHTH protein and MP, and those encoding the carbohydrate-binding domain. The former viruses encode a 5’ non-structural polyprotein followed by the capsid protein ORF whereas the latter ones encode a single polyprotein with the capsid protein domain at the N-terminus followed by the galactose-binding domain. Viruses in the family *Polycipiviridae*, the sister group of *f.0008*, encode 5’ located capsid and accessory proteins followed by the non-structural polyprotein. The galactose-binding domain is specific for a long branch within the *f.0008* family and is likely to be involved in host-specific interactions (26-28). The only other virus so far identified in *f.0008* is Lothians earthworm picorna-like virus 1 (31). The members of *Polycipiviridae* themselves were found primarily in ants and other arthropods (32) although two clades of the metatranscriptome-derived new family members use alternative genetic codes suggestive of protist hosts (1). Further inquiry into host ranges of *Polycipiviridae* and its basal *f.0008* is needed to understand evolutionary trajectories within these dynamic virus lineages and likely to split them into several more taxonomically adequate units.

With well over 4 thousand members, *Marnaviridae* is currently the largest family in *Pisuviricota* phylum (1). The characterized few viruses of this family infect diverse marine protists (33). We found that a number of marnaviruses encode an unexpected domain that is located at the C-terminus of the RdRP and shows significant structural similarity to NucS-like endonucleases (restriction endonuclease fold) (Fig 7; Dali z score 8.2). Endonucleases of different folds are also encoded by several groups of orthornaviruses, such as Influenza viruses, where this enzyme of the PD-(D/E)XK nuclease superfamily cleaves host mRNAs, snatching the cap for viral RNA synthesis (34), and nidoviruses NendoU (nidoviral uridylate-specific endoribonuclease) (35) which is involved in viral replication and evasion of host innate immunity (36, 37). The *Marnaviridae* endonuclease domain (MED) represents only the catalytic C-terminal domain of NucS-like endonucleases (‘DEK’ motif) (38), but lacks the N-terminal dimerization domain (e.g. (38)). The catalytic residues are conserved in MED, suggesting it is an active endonuclease (Fig. 7 A, B). Typically, endonucleases with this fold cleave double-stranded or single-stranded DNA (38-40) suggesting that MED could target host DNA in marnavirus-infected cells. It remains unclear whether MED is proteolytically cleaved off the *Marnaviridae* polyprotein and functions as a distinct protein or remains fused to the RdRP domain. Contigs encoding MED are scattered across the RdRP tree of *Marnaviridae* suggesting spread via HGT and/or multiple losses of the MED domain (Fig. 7 D).

**Fig. 7:**
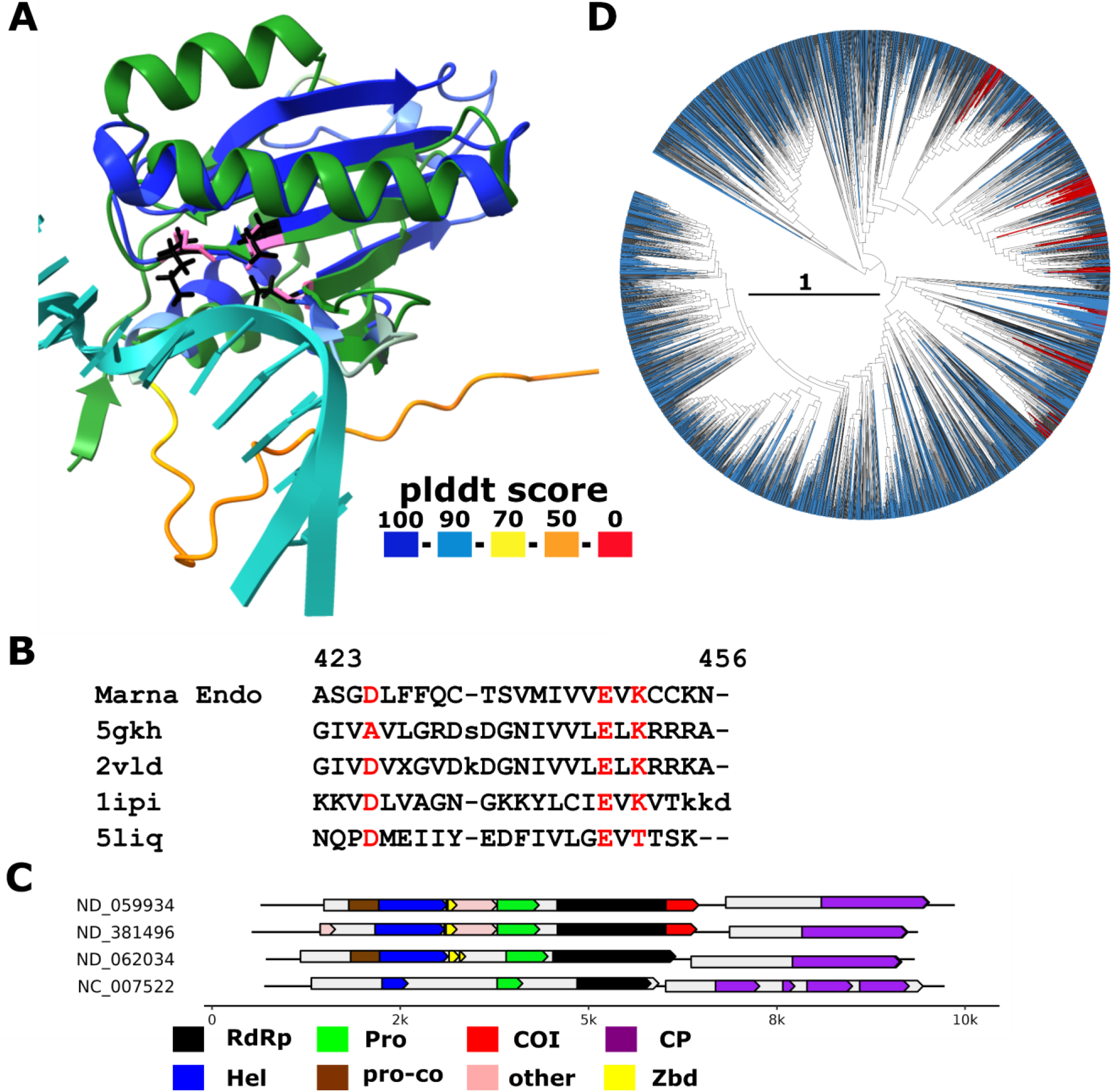
Endonuclease domain in *Marnaviridae*. **(A)** Superposition of representative *Marnaviridae* endonuclease domains (plddt score colored) with the C-terminal domain of endonuclease EndoMS (pdb 5gkh, aa 127-end, protein: green; DNA: light sea green; z-score 8.2). Catalytic residues of the nuclease are highlighted in pink for EndoMS (K181, E179 and D156A from left to right) and in black for *Marnaviridae* endonuclease (K69, E67 and D54). D156A is experimentally mutated in 5gkh to obtain the structure with uncleaved DNA. **(B)** Structure-guided alignment between *Marnaviridae* endonuclease and the four Dali top hits: endonuclease EndoMS (5gkh, Archaea, *Thermococcus kodakarensis* KOD1, z-score 8.2), NucS (2vld, Archaea, *Pyrococcus abyssi*, z-score 7.3), Holiday junction resolvase Hjc (1ipi, Archaea, *Pyrococcus furiosus*, z-score 6.7) and nicking endonuclease Nt.BspD6I (5liq, Bacteria, *Bacillus sp*., z-score 6.1). **(C)** Genome maps of representative (nearly) complete *Marnaviridae* members from (1) and ICTV exemplar (NC_007522) which either contain (top two) or lack (bottom two) the endonuclease domain. Annotations are based on profile analysis (1) and GenBank annotation (NC_007522). Protein domains: RdRp, RNA-dependent RNA polymerase, Hel, Helicase, CP, capsid protein, Pro, Protease, Pro-Co: protease cofactor_calici-como32k-like, Zbd, Zn-binding domain, COI: unannotated domain of interest (*Marnaviridae* endonuclease), other, other unannotated domain**. (D)** phylogenetic tree of *Marnaviridae* RdRps- (1); clades in which each leaf represents at least one contig that encodes an endonuclease are shown in red.

Additionally, we identified an exonuclease in the ∼450 members-strong, distinct viral family *f.0181* majority of viruses in which are likely hosted by protists that use alternative genetic codes (1). This family could not be assigned to known orders or classes in the phylum *Kitrinoviricota*. Profile annotation indicated the presence of the RdRP in these contigs but left a ∼1500 amino acid residues-long unannotated N-terminal region and a ∼300 residues-long unannotated C-terminal region in the polyprotein. Dali search revealed an DEDD superfamily exonuclease domain located at the very N-terminus (Suppl. Fig. 14, z-score: 10.6, e.g. RNase AS, a Polyadenylate-Specific Exoribonuclease of *Mycobacterium tuberculosis*). In the orthornavirus structurome, it showed structural similarities with coronavirus ExoN (z-score 8.6) which is involved in proofreading during RNA replication in these viruses possessing the largest known RNA genomes (41, 42). Given that the maximum genome size of f.0181 members is only up to 7500 nt it is unlikely that this exonuclease is involved in proofreading. Of note, the DEDDh catalytic site is modified to DEEEh in the f.0185 exonuclease domain, a variation which is not reported in literature. Moreover, structurome comparison revealed a viral methyltransferase domain within the last third of the large N-terminal region (z-score 10-11). This domain is involved in virus RNA capping during replication, a function that is widespread in diverse members of this phylum (43). The C-terminal region was identified as a SJR capsid protein by Dali search and neighboring analysis. With identification of these three functional domains, the f.0181 genome maps became less inscrutable.

The *Solemoviridae* family (*Pisuviricota*) includes four genera of plant viruses (44) and roughly two thousand newly discovered members found primarily in aquatic, soil and invertebrate metatranscriptomes (1). A fraction of solemoviruses encode an OOI with a typical α/β hydrolase fold (Fig. 8). The top Dali hits for this OOI include different types of hydrolases (for example, deacetlyases and cutinases with z-scores of 6-7), hence no clear target can be predicted. Phylogenetic distribution suggests the acquisition of this OOI in a larger clade of unclassified *Solemoviridae*, although it is not found across all genomes in this clade. Of note, we cannot detect related folds in other orthornaviruses indicating a unique acquisition of this fold or major fold-remodeling post exaptation by *Solemoviridae* members. Given that none of the known plant solemoviruses encodes this hydrolase domain and that many solemoviruses were identified in plant-less aquatic environments, it seems likely that host range of this virus family will be extended to additional host phyla.

**Fig. 8:**
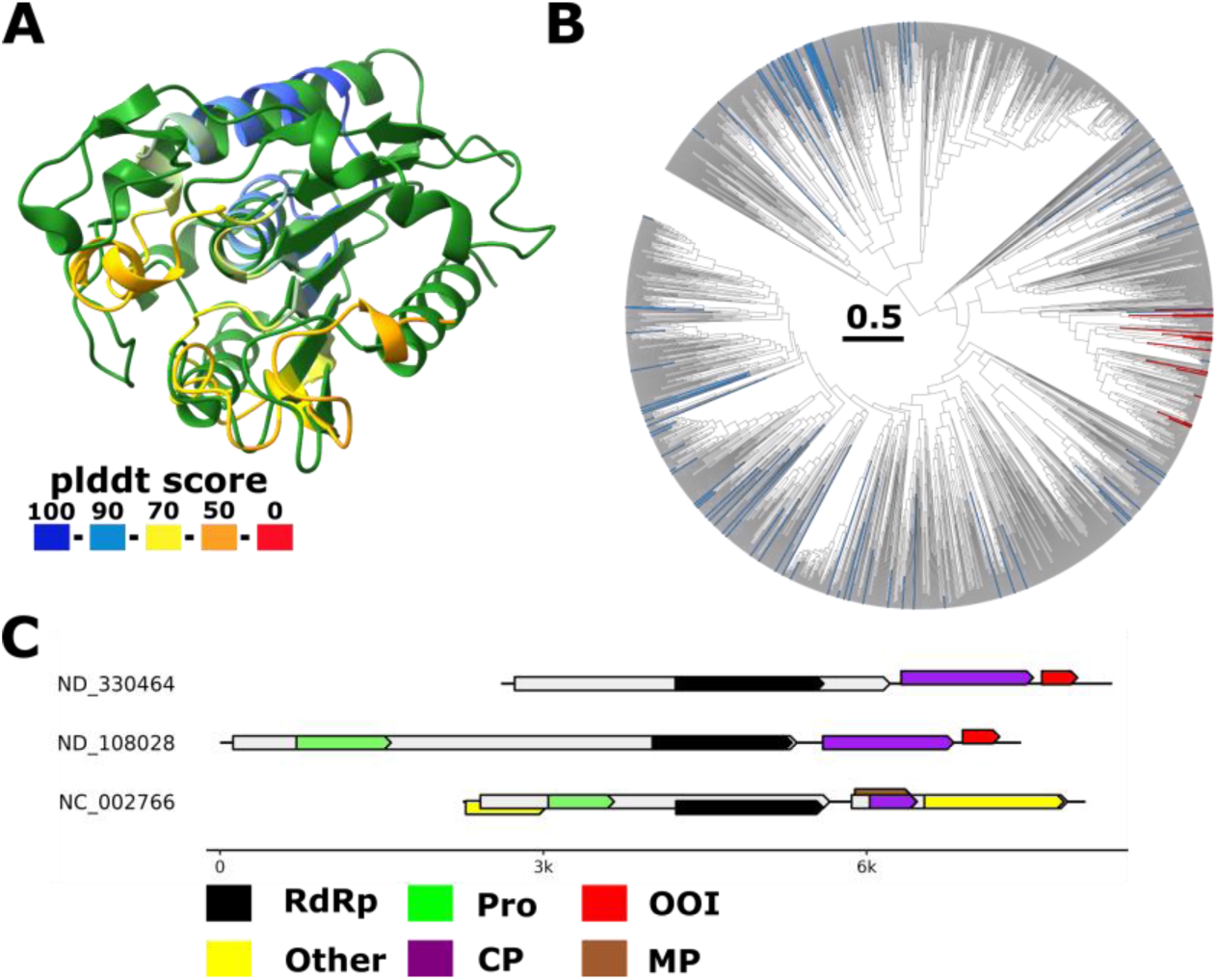
Hydrolase fold in *Solemoviridae*. **(A)** Superposition of putative hydrolase identified in *Solemoviridae* (colored by plddt score) with *Arabidopsis thaliana* SOBER1 deacetylase (pdb 6avw, green, z-score 7.0). **(B)** Phylogenetic tree with leaves representing members encoding the putative hydrolase colored red and leaves representing genomes with no coding capacity at the 3’ end for the putative hydrolase are colored blue (likely incomplete genomes). **(C)** Representative genome maps of *Solemoviridae* members. Annotations are on profile analysis (1) and GenBank annotation (NC_002766). Protein domains: RdRp, RNA-dependent RNA polymerase, CP, capsid protein, Pro, Protease, OOI, ORFan of interest, other: *Solemoviridae*-specific proteins p0 p3 and p5.

In several cases, high quality models were obtained for a COI, but there were no significant functionally characterized hits in structure comparisons. A case in point is the C-terminal domain of the *Secoviridae* (*Pisuviricota*; 57) polyprotein located immediately downstream of the RdRP (Suppl. Fig. 15). Dali search showed mainly uncharacterized proteins, the closest being the N-terminal α/β-domain of TolB (z-score of 6.8) without a known function. No significant hits (z-score >=5) were found for this COI in the orthornavirus structurome. The COI was only found in members of one *Secoviridae* genus, *Nepovirus*. Nepoviruses infect plants and are transmitted by beetles, aphids, whiteflies, leafhoppers or nematodes (45). There are no indications that this C-terminal domain is cleaved from the RdRP, so further research is needed to decipher the role of this COI during nepovirus replication and host specificity.

Similarly, an OOI encoded between the matrix and the glycoprotein in *Rhabdoviridae* (*Negarnaviricota; (46)*) members was predicted to adopt an α/βfold with 4 β-strands and 2 α-helices but without conclusive Dali hits (Suppl. Fig. 16). Orthornavirus structurome comparison showed similarity to another hypothetical protein of *Rhabdoviridae* (OA33_gp5 from NC_025382.1, a *Betapaprhavirus* member; encoded between G an L; z-score: 6.1). The function of this OOI remains unclear as is the case for other additional smaller *Rhabdoviridae* proteins that are encoded, for example, by members of the genus *Ephemerovirus* (NC 028241, Suppl. Fig. 16 C) (47).

A viral OTU (vOTU) domain, a papain-like-fold thiol protease (48), was identified in several families basal to *Deltaflexiviridae* (*Tymovirales*, *Kitrinoviricota*) family of fungal viruses (*f.0208, f.0210* and *f.0212*; Suppl.Fig. 17). Previously, a vOTU protease was identified in some members of *Tymovirales* (48), and subsequently, in *Deltaflexiviridae* (1), but not in the newly discovered basal provisional families. The present structure comparison confidently extended the spread of the vOTU protease to these families. The catalytic Cys-His dyad is conserved in all these viral proteins indicating that they are active proteases (Suppl. Fig. 17 A). In plant and animal viruses, vOTU domains function as deubiquitinylases implicated in immune evasion (49) and a similar role can be envisioned for the vOTUs of *Deltaflexiviridae* and their basal relatives.

In the same vein, we identified the SJR capsid protein in the virus family *f.0198* of the same order, *Tymovirales* (Suppl. Fig. 13). Similar capsid proteins were also found for the families *f.0194, f.0218* and *f.0217*. Based on their overall relationship to other *Tymovirales*, identification of capsid proteins in these families should have been expected, but profile comparison failed to detect these proteins due to sequence divergence.

Apart from the discovery of new domains, ORFan refinements were mainly achieved for capsid proteins, for example, for 9 distinct OOI representatives covering *Leviviricetes*, capsid proteins were identified across various families such as *Steitz*-, *Atkins*-, *Blume*-, *Solspi*-, *Fiers*-, and *Duinviridae*, but also for related putative new families in which no capsid has been so far reported (e.g. *f.0361.base-Solspi* and *f.0367.base-Atkins*). Other refinements included *Rhabdoviridae* matrix protein, phytoreovirus core-P7 dsRNA like binding domain in *Cystoviridae,* methyltransferases, E proteins or RNAi suppressor proteins in *Arteriviridae*, *Flaviviridae*, and *Dicistroviridae* and *Mymonaviridae* respectively.

## Discussion

Reliable modeling of protein structure is now available through various methods and techniques, such as AlphaFold2 and 3, RosettaFold, ESMFold, and more, and is widely applied for large-scale prediction of protein structures and functions. These analyses produced a variety of structural databases (17, 18) and many studies applying structure prediction for the exploration of evolutionary relationships among proteins and functional annotation of various genomes (17, 50, 51) (20, 52) (52). To our knowledge, however, there are currently no curated large-scale databases available for virus-encoded proteins although Nomburg *et al.* presented a study on predicted virus protein structures when drafting this manuscript (19). For example, as of June 10^th^ 2024, EBI AlphaFold repository dismissed viral proteins until computational polyprotein processing would improve (https://alphafold.ebi.ac.uk/ (18)). Here, we sought to start closing this gap by modeling the structures of the proteins from 498 orthornavirus families using AlphaFold2. The majority of unannotated domains and putative ORFs yielded low-quality models which could be expected given that unannotated ORFs and protein regions are a highly heterogeneous set that includes variable sequences, linkers and intrinsically disordered protein-protein interaction interfaces. Furthermore, some of the smaller ORFans could have been acquired recently or evolved *de novo*, are virus-specific and are unlikely to have a large footprint in the databases used for model prediction. Nevertheless, we identified a large set of high-quality models for both CUDs and ORFans many of which could be linked to structures from proteins with known functions.

The spread of almost all of the predicted new structures and functions was limited to a single virus family, and typically, only to a subset of its members. Thus, the general conclusion from this work is that the current catalogue of widespread protein domains of orthornaviruses is effectively complete, even for novel groups known only from metatranscriptome mining. The dark matter of the orthornavirus pan-proteome consists mostly of all-helical and intrinsically disordered domains and proteins that are not readily amenable to structure modeling and comparison. This conclusion on the near saturation of the orthornavirus domain repertoire contrasts the ever-expanding diversity of the RdRPs that shows no signs of saturation (1-4, 53). This difference reflects the strong constraints on the size of RNA genomes that limit the potential for new gene capture, in a sharp contrast to viruses with large DNA genomes.

The general conclusion on the limited domain repertoire of orthornaviruses notwithstanding, the globular domains that we identified here do show some trends. Specifically, some of these are predicted nucleic acid-binding domains and nucleases that could be involved in viral interference with host-specific immune systems. Typically, the ORFans with predicted new structures and functions are not found in a single viral clade but rather are scattered over the evolutionary trees of the respective viral families. This pattern is likely to reflect dynamic evolution of these relatively recently captured genes involving multiple HGT events as well as gene loss. Generally, uncharacterized regions of known viral (poly)proteins in which globular domains were predicted span broader taxonomic ranges of viruses than ORFans (48). This difference seems plausible because acquisition or *de novo* emergence of a new small ORF while maintaining a replication competent virus appears to be more likely than insertion of a new domain inside a large virus protein.

Somewhat serendipitously, during this structural analysis of the uncharacterized proteome of orthornaviruses, we identified a notable case of exaptation of a host enzyme, NMP kinase, for dsRNA-binding function. This protein is widely spread across diverse orthornaviral families suggestive of an important role(s) in virion morphogenesis but, possibly, also in suppression of host immunity. Exaptation of enzymes for structural roles seems to be a common theme in the evolution of large viruses with dsDNA genomes, such as poxviruses (54), but is less frequent in orthornaviruses (55) and other viruses with small genomes. More generally, exaptation of the host- and virus-encoded proteins is a leading trend in the evolution of viruses (56).

The expansion of the RNA virosphere via metatranscriptome mining is ongoing at an accelerating pace, and novel viruses can be confidently expected to emerge for many years to come, at least, at the levels of family and below. Thus, the pangenome and structurome of orthornaviruses constructed in this work, along with advancing tools for protein structure modeling and comparison, should be helpful to researchers investigating the structural and functional diversity of the RNA virosphere.

## Conclusions

The current analysis of the orthornavirus pan-proteome and its dark matter using structure prediction and comparison methods suggests that all broadly conserved proteins and domains encoded by viruses of this kingdom are already known. The proteomic dark matter seems to consist largely of disordered and all α-helical proteins that cannot be readily assigned a specific function. It appears likely that these domains mediate various interactions between viral proteins and between viral and host proteins. Nevertheless, we identified a substantial number of globular domains that have not been reported previously. These domains are primarily encoded by ORFans, that are present only in narrow groups of orthornaviruses or are scattered across several such groups. With the accelerating discovery of orthornaviruses in metatranscriptomes, protein structure modeling and analysis is now the approach of choice for the characterization of lineage-specific viral proteins and domains. The orthornavirus structurome constructed in this work can be expected to facilitate such studies.

## Materials and Methods

### ORF and domain identification

Virus contigs and protein annotations were retrieved from the data deposited by Neri *et al.* (1) (<u>DOI:10.5281/zenodo.7368133</u>), named EMRV set (environmental metatranscriptome RNA viruses) hereafter. ORF boundaries were identified by running emboss getorf (minimal number of nucleotides: 150, stop-stop to allow for the identification of incomplete ORFs in incomplete genomes) with the standard genetic code or the code indicated in (1). Putative start codons were identified within the extracted ORFs and assumed to be ORF starts unless the ORF was located at the very 5’ end of the contig (likely incomplete assembly) or overlapped with an annotation from (1) (see below, possibility of programmed frame-shift). Annotations from (1) were mapped back to the ORFs. An ORF (and its protein sequence) was called fully annotated if it contained no continuous unannotated stretches of 60 aa or more. Otherwise, the annotated part was extracted as domain, retrieving in total 647,383 annotated ORFs and domains.

To identify conserved unannotated domains (CUD) and ORFs, a pipeline was run as follows: At the first step, all proteins larger than 200 aa with partial or no annotation (297,411 proteins, with at least one stretch of at least 60 aa unannotated) were clustered and aligned using the snakemake pipeline (57) (Suppl. file ‘protein-clustering-diamond-mcl_full.smk’ ran with snakemake --config input=proteins.faa precluster_min_seq_id=0.95 diamond_min_seq_id=0.0 min_aln_cov=0.9 mcl_inflation=2.5 -j 16 -s protein-clustering-diamond-mcl_full.smk). In detail, sequences were pre-clustered with mmseqs2 (mmseqs easy-linclust --kmer-per-seq 100 -c 1.0 --cluster-mode 2 --cov-mode 1 --min-seq-id 0.95) (58) and representatives were searched against each other using diamond (diamond blastp -e 1e-3 --very-sensitive --id 0.0) (59). Pairs with at least 90% coverage were identified and clustered with mcxload (mcxload -abc {input} --stream-mirror --stream-neg-log10 -stream-tf ’ceil(300)’ -o {output[0]} -write-tab {output[1]}) (60) and mcl (mcl {input[0]} -use-tab {input[1]} -o {output} -te {threads} -I 2.5) (60). Protein clusters including 5 or more sequences were kept to focus on domains from abundant proteins. Clusters were aligned with mafft (61) and written to a file with seqkit (62) (mafft --quiet --anysymbol --thread 4 --auto {input} | seqkit seq -w 0 > {output}), resulting in 2,410 alignments spanning 31,041 sequences. These alignments were used to conduct a first iteration of hhblits (63) to identify known protein domains. First, fasta alignments were converted to a3m alignments with hhconsensus (-M 50) (63) and searched against the following databases: pdb70 (64), pfama (65), scope70 (66, 67), ECOD (ECOD_F70_20220613) (68) and nvpc (1) (hhblits -cpu 1 -norealign -n 1 -p 0.9 -z 0 -Z 5000 -b 0 -B 5000 -i {input} -o {output} -d {pdb70} -d {pfama} - d {scope70} -d {ECOD} -d {nvpc}). Unannotated stretches of at least 60 aa in the alignment were extracted if not overlapping with an annotation (90% probability) by more than 10% (Suppl. File get_uncovered_coord_first_round.py and snakemake files snakemake_interdomain_p1.sh and snakefile_interdomain_p1.smk). The extracted stretches of unannotated alignments were run with hhblits against the same databases with the same parameters and processed as in the first round to retrieve final unannotated alignment stretches. This procedure resulted in 2,831 unannotated alignment stretches spanning 34,710 domains (31,247 larger than 60 aa which were kept as CUDs (conserved unannotated domains)).

To identify abundant unannotated ORFans between 60 and 200 aa in size, only ORFs between start and stop codons were considered (no programmed frameshifts or incomplete ORFs at the 5’ end of the genome lacking the start codon included). In the EMRV set, 1,363,871 such unannotated ORFs wsere detected including those located on either the forward or the reverse strand and those nested with other ORFs. ORFans were clustered together with CUDs using mmseqs2 (-min-seq-ident 0.4 and -c 0.85). Only clusters with at least one CUD or at least 5 ORFans were considered, resulting in 59,815 clusters spanning 655,787 ORFans and 13,099 clusters consisting of CUDs including 325 clusters that included 1052 ORFans together with CUDs. Representatives from these clusters were kept for protein structure prediction.

### Processing of ICTV exemplars

To obtain a reference set of functional domains represented in each virus family, reference virus genomes were downloaded from ICTV exemplars for all Orthornavirae (https://ictv.global/vmr, VMR_MSL38_v2, Dec. 1^st^ 2023). GenBank files of viral genome were retrieved from NCBI for all ICTV approved virus families in the EMRV set (98 virus families). Proteins and domains were extracted as annotated within the genome GenBank file. To account for a more fine-grained annotation of individual proteins, GenBank files for individual proteins were retrieved whenever the protein in the genome GenBank file was flagged as polyprotein or was at least 500 amino acids long. Again, individual domains were isolated and mapped to the corresponding virus family (32,648 domains in total). To compare the GenBank annotation with our profile-based annotation, isolated ICTV exemplar domains were run against the viral profile database from (1) (nvpc db) using hhsearch with default settings. Hits with 95% or higher probability were harmonized by taking the most prevalent function. All domains were then clustered using mmseqs2 (min-seq-id 0.4, coverage 0.85; 9245 clusters) and profile hits were harmonized across all members of a cluster. The most frequent function was then assigned to all members of a cluster. Clusters that lacked at least one highly significant profile hit were inspected for GenBank annotation of the cluster members (4,697 sequences across 3,163 clusters). Those GenBank annotations were harmonized by keyword (e.g. ‘hypothetical’, ‘unknown’ and ‘putative protein’ were all denoted ‘hypothetical’) and the most frequent label was assigned to the cluster. The clusters were then assigned to the respective virus families and profile annotation coverage per virus family was calculated (Suppl. Fig 2). This procedure was performed for clusters harboring either all domains or domains encoded on the RdRp-encoding segment for viruses with segmented genomes.

### Building pangenomes of orthornavirus families

Annotated domains from the EMRV set were mapped to the ORFs and assigned to their respective virus families. Nucleotide and amino acid positions were orientated such that the RdRp encoding ORF is on the forward strand on each genome. Next, CUDs were mapped to the respective genomes. As CUDs were retrieved by a slightly different method than the annotation in (1), 10 aa overlaps between CUDs and annotated domains were permitted.

### Protein structure prediction, analysis, comparison and visualization

Protein structures were modeled using AlphaFold2 (version 2.3.1, default parameters) for all proteins and domains of orthornaviruses (annotated, unannotated and ICTV exemplars) except ORFans (all Prodigal annotated viral proteins from (1) were added to the uniref90 database to improve alignments) on the high-performance biowulf cluster at NIH. ORFans were modeled using ESM-fold (version 2.0.0) because of the prohibitive computational cost of modeling such a large number of sequences with AlphaFold 2. Model quality was assessed by the plddt score. Protein structures were clustered with foldseek (-c 0.8) (69). Protein structures of selected representatives of CUD and ORFan clusters with plddt score of 70 or higher were searched against a local version of pdb70 using Dali (23). Protein secondary structure was assessed by psique (24) for all CUDs and ORFan structures and based on the Dali search files for representative structures ran against pdb (an individual helix was called if at least 10 consecutive amino acids were assigned as helix and a beta-strand was called if at least three consecutive amino acids were assigned as beta-strand member). Protein structures were visualized with Chimera X (version 1.3) (70).

### The orthornavirus ‘structurome’

To obtain the orthornavirus ‘structurome’, one representative protein of each functional tag per virus family was taken from the Riboviria pangenome and from ICTV exemplar domains lacking confident annotation, the structures of these representatives were predicted as described above and combined with representative structures of OOIs and COIs. All 7,996 structures were pre-clustered with foldseek (-c 0.8) and 5,528 representatives were ran all-vs-all using Dali (default parameters). Clusters were identified using an iterative process in which, first, all closest related pairs across the all-vs-all matrix were identified (based on their z-score, minimal z-score here 7, ‘z7 cluster’) as preliminary clusters. Then, the next closest related pairs were identified. Whenever the members of a pair belonged different clusters, such clusters were merged if the mean inter-cluster z-score was 7 or higher. In case only one member was part of a cluster, the second one was added if the mean z-score to all members in a cluster was 7 or higher. In case none of the members of a pair belonged to a cluster, a new preliminary cluster was created. The iterations stopped when no new members could be added, and no clusters could be merged.

A structure similarity network was constructed based on Dali z-scores of the all-vs-all comparison to visualize the structure relationships. All structures present in a z7 cluster were considered. Structures outside of such clusters with a z-score of 7 or higher to any structure within a cluster were also included. Each structure was a node and each edge linking two nodes was weighted by the z-score. Node and edge tables were loaded into Gephi (71) (version 0.10.1) and arranged by running the openOrd algorithm with default parameters (Stages and their percentages: liquid (25%), expansion (25%), cooldown (25%), crunch (10%) and simmer (15%), Other parameters: Edge cut: 0.8, Nin threads: 5, Num iterations: 750, fixed time: 0.2, random seed: 2644300876718762555) followed by a round noverlap (speed: 3, ratio: 1.2, margin:2).

### Provisional annotation of CUDs and ORFans

All initially extracted 31,247 CUDs were run against a blast database of all annotated domains from the EMRV set as well as all ICTV exemplar domains using PSI-BLAST (72) (default parameters, eval 0.0001). Annotations were mapped on the 13,085 CUD mmseqs clusters and the cluster representative provisionally labeled if at least 50 % of the members produced a hit; the hits were then harmonized. If no harmonization was possible, the hit with the lowest e-value was taken. Through this procedure, 37% of the 13,085 CUD mmseqs representatives were annotated.

ORFans and CUDs were then clustered together with ICTV exemplar domains using mmseqs2 (min-seq-ident 0.4, coverage 0.85) and clusters were aligned with mafft (61) (default parameters). Aligned clusters were searched against various databases ((ECOD (68), scope70 (66, 67), pfam (65), ncbi CD (73), nvpc (1), virusDB2020 (74)) using hhblits (63) (default parameters). Clusters that produced hits with a probability of 90% or higher were provisionally annotated. The ICTV exemplar domain annotation was considered for a cluster whenever such a domain was present and no hhblits hit with a probability of 90% or higher was obtained. About 1% of the clusters were provisionally annotated.

### Identification of CUDs of interest (COI)

All 13,000 CUD models were filtered by their mean plddt score (70 or higher) and secondary structure. To be kept, a CUD had to encompass at least 3 helices or one helix and one beta-strand or more than 2 beta strands based on the Dali secondary structure assignment, where at least 10 consecutive ‘helix’ assigned amino acids counted as a helix and at least 3 consecutive ‘beta-strand’ assigned amino acids counted as a beta-strand. This procedure excludes single helices, helix-turn-helix folds, beta-hairpins and disordered proteins, resulting in 830 CUDs of interest (COI). Then, COIs were compared to the N- and C-terminal annotated domains in the same genome to see whether the COI might be an extension of the existing annotation. To determine whether a COI was annotated already in a related protein, genomic neighborhood analysis was performed: similar proteins with and without the COI annotation were aligned using mafft (61) (default parameters) and annotated domains and COI positions of each included protein were mapped to the alignment. If the COI region overlapped by 50% or more with any annotated domain of another protein in the alignment, the provisional COI PSI-BLAST annotation (if any, see above) was compared with that of the profile-annotated domain. If there was no conflict, the COI was called as ‘resolved by neighborhood’. In case of a conflict, the Dali structure comparison results for the given COI were manually inspected. If there was no overlap by at least 50%, the COI was kept for manual inspection. Only COIs with a top Dali z-score of 4 or higher were inspected. In addition, all-helical proteins were only inspected if the top z-score was above 10 given that all-helical proteins tend to produce inconclusive results because helices in many different types of proteins are hit. None of the all-helical COIs with a top z-score above 10 produced a conclusive hit. Furthermore, randomly checked all-helical COIs with lower top z-score could not be assigned to specific folds. The remaining COI structures with a Dali top z-score of 4 or higher were inspected with respect to i) the respective Dali pdb hits, ii) their position in the genomes of a virus family, iii) their relationship to other structures in the structurome and iv) their provisional hhblits annotation, if any.

### Identification of ORFans of interest (OOI)

The modeled ORFans were clustered with foldseek (-c 0.8) and inspected by their plddt score. ORFans present in a cluster with a plddt score of 70 or higher were further analyzed (5,944 / 59,185 ORFan structures). Structure representatives were run against a local version of pdb70 using Dali. As for CUDs, structures with ‘simple’ folds were excluded (see above), leaving about 1,600 ORFans of interest (OOIs). As for COIs, OOIs were manually inspected whenever they were not all-helical. The remaining COI structures with a Dali top z-score of 6 or higher were inspected with respect to i) the respective Dali pdb hits, ii) their position in the genomes of a virus family, iii) their relationship to other structures in the structurome and iv) their provisional hhblits annotation, if any.

### Detection of intrinsically disordered regions

The fraction of intrinsically disordered regions in a protein was analyzed using a local version of MobiDB-lite (version 3.10.0) (75).

### Phylogenetic distribution of OOIs and COIs

For each virus family, representative OOIs or COIs of the same function were aligned with their foldseek and mmseqs2 cluster members using muscle ((76), version 5.1, default parameters). Aligned sequences were searched using PSI-BLAST (77) (version 2.15) (psiblast -in_msa ip -threshold 9 -db db - evalue 0.01 -out op -num_threads 8 -max_target_seqs 50000 -outfmt 6) against all extracted ORFs (annotated or not) from the EMRV set. All regions covered were extracted, realigned together with the initial sequences and ran again against all ORFs using PSI-BLAST with the same parameters. The final sequence sets were mapped back to the respective virus family phylogeny obtained from (1). Each leaf covering at least one genome containing the respective OOI or COI was called positive. The leaves in each tree represent clusters of RdRp core sequences at 95% sequence identity (1). Trees were visualized using iTol (v6) (78).

### Visualization of viral genomes

Representative viral genome maps were generated with R using the libraries ggplot2 and gggenomes (https://github.com/thackl/gggenomes).

### Structure-guided alignment of core-P7 RBD domains and cellular kinases and tree construction

Representative structures of viral core-P7 RBD domains and related cellular kinases were aligned using the FoldMason web server (26). The alignment was trimmed to remove columns with more than 35% gaps, and a phylogenetic tree was built using IQtree2 with the implemented modelfinder (-m MFP (79)) and ultrafast bootstrapping (-B 10000 (80, 81)).

### Sampling of unique domains across *Orthornavirae*

Increasing numbers of clusters of similar genomes (based on 90% RdRP aa identity, each cluster corresponding to a leaf in the virus family phylogeny) were sampled randomly across all *Orthornavirae* (step size: 50 clusters; till all clusters were sampled) and the number of distinct domains was extracted. This procedure was bootstrapped 30 times and the mean, minimal and maximal number of unique domains plotted as function of the number of sampled clusters.

### Transmembrane domain prediction

TMHMM 2.0 (82, 83) was used to predict transmembrane domains in CUDs and ORFans.

### Defining the ‘core’ set of unique domains per virus family

Distribution of associated contig lengths was obtained for each virus family; contigs of length of at least 2/3rd of the 75th percentile we operationally classified as “near full-length”. For families with at least 10 near full-length contigs, the full set of identified structural domains was identified along with their frequencies. All domains were ranked by their frequencies; for domains, present in at least 50% of the near full-length contigs, the “frequency gap” (difference between its frequency and that of the next-ranked domain) was calculated; the domain with the highest frequency gap was defined as the last core domain.

## Availability of data and materials

This work is based on the analysis of genomes publicly available in GenBank. All other data generated by this analysis are contained in the Supplementary Material or publicly available at <u>zenodo (</u>10.5281/zenodo.13770615)]

## Supplementary files

Pangenome pickle file

ICTV exemplar pangenome pickle file

Fasta files: annotated domains, un-annotated ones, ORFans

Table z7 cluster function

## Author contributions

P.M. and E.V.K. conceptualized the project; P.M., A.P. C., H.S., A.B., U.N. and Y.I.W. developed the methodology and collected the data; P.M. performed research; P.M., M.K., V.V.D. and E.V.K. analyzed the data; P.M. and E.V.K. wrote the manuscript which was read, edited and approved by all authors.

## Acknowledgements

P.M., H.S., Y.W. and E.V.K. are supported by the Intramural Research Program of the National Institutes of Health (National Library of Medicine). This work utilized the computational resources of the NIH HPC Biowulf cluster (http://hpc.nih.gov). V.V.D. research was supported in part by an appointment to the National Center for Biotechnology Information Scientific Visitors Program administered by the Oak Ridge Institute for Science and Education (ORISE) through an interagency agreement between the U.S. Department of Energy (DOE) and the National Institute of Health. ORISE is managed by ORAU under DOE contract number DE-SC0014664. All opinions expressed in this paper are the author’s and do not necessarily reflect the policies and views of NIH, NCBI, DOE, or ORAU/ORISE. A.B. is supported by a post-doctoral fellowship from Foundation pour la Recherche Mèdicale (grant number SPF202110014092).

## Supplementary figures

**Suppl. Fig.1: Workflow schematic**

Workflow and numbers for the identification and extraction of annotated and un-annotated proteins and domains with Part 1 and 3 covering the domains and part 2 the putative ORFs <200 aa.

**Suppl. Fig.2: ORFan and CUD clustering**

ORFans and conserved unannotated domains (CUDs) were clustered with mmseqs (ident. 0.4, coverage 0.85) and numbers of leaves in the RdRp-based phylogenetic trees represented by each cluster identified. **(A)** Overall cluster size distribution. **(B)** Leaves represented per mmseqs cluster. Clusters covering at least three leaves were further considered.

**Suppl. Fig.3: Profile annotation coverage across ICTV exemplar protein clusters**

ICTV exemplar proteins and domains were clustered with mmseqs2 (ident 0.4, coverage 0.85) and searched against nvpc db by hhsearch. **(A)** Percentage of clusters of ICTV exemplar domains and proteins per virus family confidently covered by profile-based annotation. Left: for proteins on all viral segments. Right: only considering RdRp-encoding segments. **(B)** Functional annotation from genbank file for ICTV exemplar domains and proteins not covered by profile-based annotation. **(C)** ICTV exemplar protein and domain clusters not covered by profile-based annotation per virus family.

**Suppl. Fig.4: Characteristics of ORFan structures**

**(A)** 6117cluster representative ORFans were modeled with AlphaFold2. Plddt score vs sequence length is shown. **(B)** Same as A but colored by fraction of protein length identified as disordered by MobiLite. **(C)** Foldseek (cov. 0.8) cluster size distribution of the 6177 ORFan structures. **(D)** Secondary structure assignment based on Dali from 412 representative ORFans which have a plddt score >=70.

**Supp. Fig. 5: Characteristics of CUD structures**

**(A)** 13,085 cluster representative CUDs were modeled with AlphaFold2. Plddt score vs sequence length is shown. **(B)** Same as A but colored by fraction of protein length identified as disordered by MobiLite. **(C)** Foldseek (cov. 0.8) cluster size distribution of the 13,085 CUD structures. **(D)** Secondary structure assignment based on Dali for representative CUDs with plddt score >=70.

**Suppl. Fig. 6: Phylogenetic distribution of OOI and COI structure clusters**

Number of clusters are shown and the phylogenetic distribution across four taxonomic ranks (phylum, class, order and family) of the cluster members per cluster. (**A) and (C)**: For ORFans of Interest (OOI), **(B) and (D):** For CUDs of interest (COI). A and B show all clusters, **(C) and (D)**: those clusters whose members are found in more than one unique taxon of the given taxonomic ranks. Colors: blue, orange and purple: one, two or three or more taxon(s) per taxonomic rank for all cluster members, respectively.

**Supp. Fig. 7: Structurome clustering**

**(A)** Cluster size distribution after foldseek (cov. 0.8) clustering of 6,335 structures (852 COIs, 146 OOIs, 2,421 annotated domains and 3,000 ICTV exemplar domains). **(B)** Cluster size after Dali all-vs-all run of 4,022 cluster representatives from A. Iterative cluster identification in which each member has an average Dali z-score of 7 or higher to each other structure in the cluster when added. Domains indicated for largest clusters. RdRp: RNA-dependent RNA polymerase, Cap MTase: viral capping methyltransferase, PRO-Chy: chymotrypsin-like protease, SJR: single jelly-roll fold, all helical: all alpha-helical folds without clear functional assignment across all members of the cluster.

**Suppl. Fig 8: Orthornaviral structurome as a structure-structure network**

Structure-structure network presentation of the Dali all-vs-all run of 4,022 structure representatives. Each node represents a structure connected to another node by an edge which is weighted by the Dali z-score. Edges are present between structures of the same Dali z-score of 7 cluster or between any node not present in such a cluster and a member within a cluster if the z-score was 7 or higher. Functional domains, COIs and OOIs are colored.

**Suppl. Fig. 9: Beta-barrel folds found in *f.0145***

Two tandem OOIs with distinct beta-barrel fold found in f.0145 members (plddt score colored) superimposed with their top Dali hits **(A)** Pilz domain in *E.coli* YcgR (pdb 5y6f, green, aa 125-243, z-score 5.5)and **(B)** Pseudomonas aeruginosa protein HCP3 (light green, z-score 5.6). **(C)** Phylogenetic tree of f.0145 with leaves covering genomes encoding for both beta-barrel-fold OOIs (red) or only the Pilz-like OOI (orange). **(D)** Representative genome maps of f.0145 members. Annotations based on profile annotation (Neri *et al*.). Protein domains: RdRp: RNA-dependent RNA polymerase, CP: capsid protein, OOI: ORFan of interest (1: Pilz-like, 2: HCP3-like).

**Suppl. Fig. 10: Structure-guided alignment of viral core-p7 RBD-like folds and related cellular kinases**

**(A)** Structure guided alignment using FoldMason web server (26) from viral core-p7 RBD-like structures from various viral families with related kinases. See details on cellular kinases in legend of Fig. 4. Position of Walker A and Walker B indicated. Note, that Shikimate kinases (1shk, 4y0a and 3nwj) have an atypical Walker B motif of *hhh*GGG with *h* being hydrophobic residues (black box).

**Suppl. Fig. 11: Viral core-p7 RBD-like folds from various viral families and genome organization**

**(A-H)** AlphaFold2 predictions of core-p7 RBD-like domains from *Picobrinaviridae* (A,B), *Flaviviridae* (C), *Endornaviridae* (D), *Phytoreovirus* p7 with enlarged RBD domain (E), *<u>Ghabrivirales</u>* families *Chrysoviridae* (F), *f.0269* (G) and *f.0285* (H). Walker A motif (black) and former Walker B motif (gray) indicated, otherwise plddt-scorer colored. **(I)** Representative genome maps of genomes encoding for a core-p7-RBD-like domain.

**Suppl. Fig. 12: VSR-like RNA binding domain in f.0092.base-Permutotetra**

**(A)** Superposition of dsRNA binding domain found in *f.0092.base*-*Permutotetraviridae* (plddt colored, dsRNA binding domain aa 140-226) with a RNA-binding domain (dsRBD) of human adenosine-deaminase ADAR1 (2mdr, green; z-score 8.4). **(B)** Phylogenetic tree for *f.0092.base-Permutotetra* as from Neri *et. al.* Branches colored in red in case a COI is present in at least one contig represented by the leaf or in blue in case all contigs represented by the leaf might be incomplete and therefore not showing the COI. In this case: if there are less than 60 aa unannotated after RdRp annotation the contig might be incomplete/ stop codon due to sequencing error. **(C)** Genome maps of representatives encoding the dsRNA binding domain. Protein domains: RdRp: RNA-dependent RNA polymerase, CP: Capsid, coi: domain of interest.

**Suppl. Fig. 13: Galactose binding domain in *f.0008.base-Polycipi***

**(A)** Superposition of representative COI (plddt score colored) with a family 32 carbohydrate binding module (CBM) from Clostridium perfringens NanJ (2v72, green, z-score: 15.4). **(B)** Genome maps of representatives encoding the carbohydrate binding module. Protein domains: RdRp: RNA-dependent RNA polymerase, Hel: Helicase, Pro: Protease, CP: Capsid, COI: conserved unannotated domain of interest, other: other un-annotated domain. **(C)** Phylogenetic tree for *f.0008.base-Polycipi* as from Neri *et. al*. Branches colored in red in case a COI is present in at least one contig represented by the leaf or in blue in case all contigs represented by the leaf might be incomplete and therefore not showing the COI. In this case: If no N-terminal capsid protein annotation could be found within any contig represented by the respective leaf.

**Suppl. Fig. 14: Exonuclease & methyltransferase in *f.0181***

**(A)** Superposition of representative COI (plddt score colored) with RNase AS, an exonuclease from Mycobacterium tuberculosis (green, z-score 10.6) and viral capping methyltransferase from family *f.0176* (AlphaFold2 modeled annotated domain, cyan, z-score in structurome: 11.0). **(B)** Catalytic residues of the exonuclease domain for COI (black, D14, E16, E91, E125, and H120) and RNase AS (4okk, magenta, D6, E8, D95, D145, H140 (DEDDh)). **(C)** Representative genome map of *f.0181* member. Protein domains: RdRp: RNA-dependent RNA polymerase, Coi: un-annotated domain of interest with ExoNuc (exonuclease) and MetTrans (viral capping methyltransferase), other: other COI with refinement of capsid protein (CP). **(D)** Phylogenetic distribution of full COI across f.0181 (red). Blue branches indicate putative incomplete genomes with less than 60 aa N-terminal of annotated RdRp domain.

**Suppl. Fig. 15: Secovirus C-terminal domain of unknown function**

**(A)** Superposition of representative domain (plddt score colored) with E.coli TolB (2ivz, green, aa 34-162, missing the unstructured N-term and C-terminal beta-propeller; z-score 6.8). **(B)** Genome maps of representative *Secoviridae* contigs (RdRp segment). Protein domains: RdRp: RNA-dependent RNA polymerase, Hel: Helicase, Pro: Protease, Pro-Co: pro_cofactor_calici-como32k-like, coi: un-annotated domain of interest, other: other un-annotated domain. **(C)** *Secoviridae* phylogenetic tree as in Neri *et al.* Branches colored in red in case a COI is present in at least one contig represented by the leaf or in blue in case all contigs represented by the leaf might be incomplete and therefore not showing the COI. In this case: if there are less than 33 aa unannotated after RdRp annotation the contig might be incomplete/ stop codon due to sequencing error.

**Suppl. Fig. 16: alpha-beta fold OOI found in *Rhabdoviridae***

**(A)** Superposition of alpha-beta-fold OOI found in Rhabdoviridae members (plddt score colored) and an hypothetical ORF from a member of the genus *Betapaprhavirus* (NC_025382.1, OA33_gp5, located between G and L protein, green, z-score in structurome: 6.1). **(B)** Phylogenetic tree of Rhabdoviridae with red leaves representing genomes encoding for the OOI (red) and blue ones representing genomes without coding capacity between the M and G protein. **(C)** Representative genome maps of *Rhabdoviridae* members. Annotations based on profile annotation (Neri *et al*.) and gb annotation (NC_020803 and NC 028241). Protein domains: RdRp: RNA-dependent RNA polymerase, N: Nucleocapsid, P: Phosphoprotein, M: Matrix protein, G: Glycoprotein, MetTransf: viral capping Methyltransferase, *Ephemerovirus* specific proteins: ORFs found specifically in the genus *Ephemerovirus.* OOI: ORFan of interest.

**Suppl. Fig. 17: vOTU in *Deltaflexiviridae* and related families**

**(A)** Superposition *of f.0212.base-Deltaflexi* viral OTU domain representative (plddt score colored, aa 1-121) with viral OTU proteinase from Turnip yellow mosaic virus (pdb id 4a5u, green, aa 732-879 of the polyprotein; z-score 6.43). Catalytic dyad is shown in black (C2, H95) and pink (C783, H869), respectively. **(B)** Genome map of representatives of the family *f.0212.base-Deltaflexi*. Protein domains: RdRp: RNA-dependent RNA polymerase, Hel: Helicase, put. Pro: putative Protease, MetTrans: viral methyltransferase, uoi: un-annotated domain of interest, other: other un-annotated domain.

**Suppl. Fig. 18: Capsid protein in *f.0198***

**(A)** Superposition of capsid protein found in *f.0198.base-Tymo* (plddt score colored) with tymovirus capsid (pdb 1ddl, green, z-score 10.9). **(B)** Phylogenetic distribution of leaves representing genomes encoding for the capsid (red) or lacking coding capacity after the polyprotein (blue, likely incomplete genomes). **(C)** Representative genome maps for f.*0198* members with and without OOI. Protein domains: RdRp: RNA-dependent RNA polymerase, Hel: Helicase, Prot: Protease, MetTrans: viral capping Methyltransferase, OOI: ORFan of interest.

**Suppl. Fig. 19: Distinct domains per virus family**

(A) The number of distinct domains per virus family by median genome length. (B) As in A) but for the core set of distinct domains genomes.

