## Supplementary figures for "The protein structurome of *Orthornavirae* and its dark matter"

**Suppl. Fig.1: Workflow schematic**

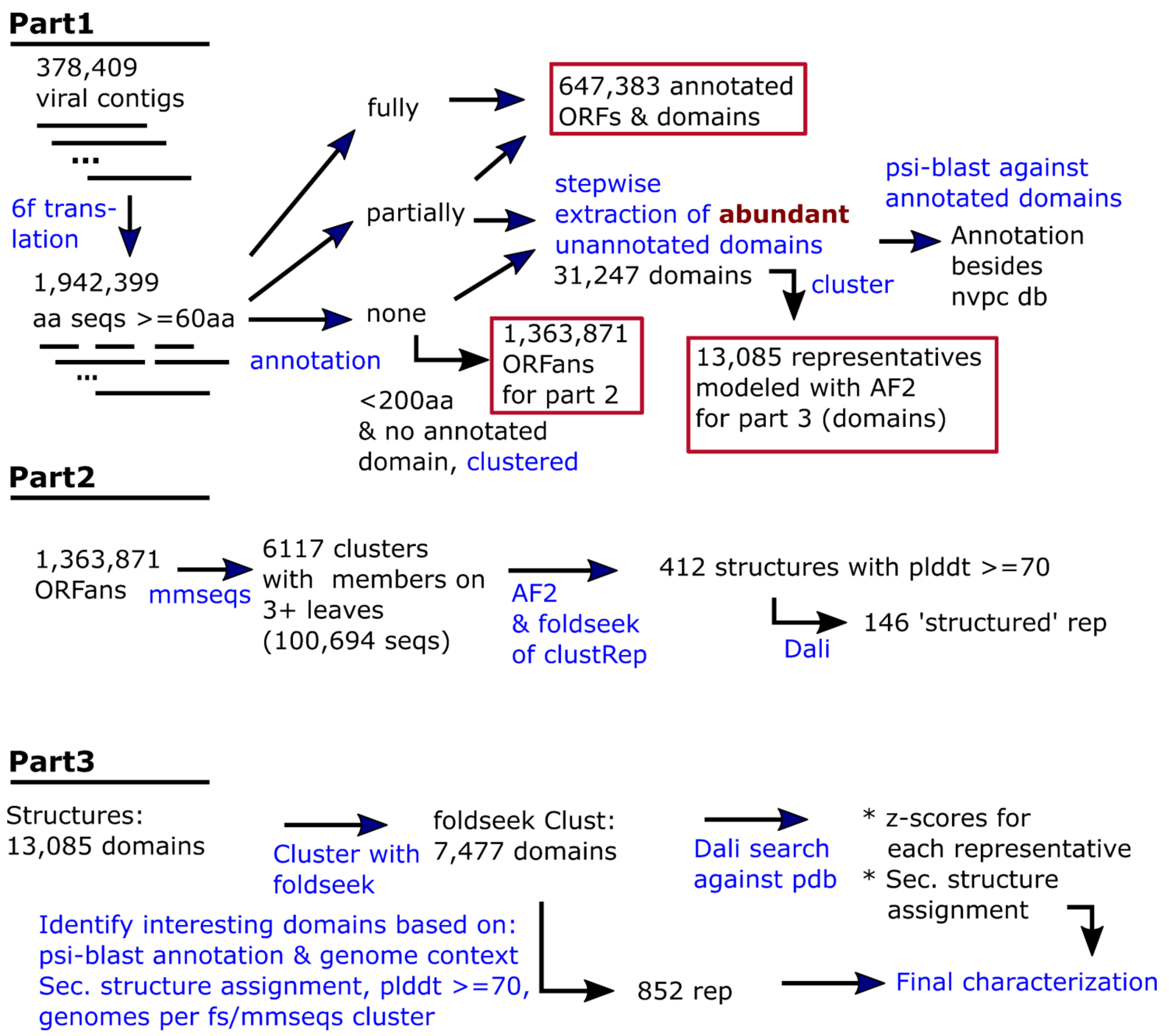

**Suppl. Fig.2: ORFan and CUD clustering**

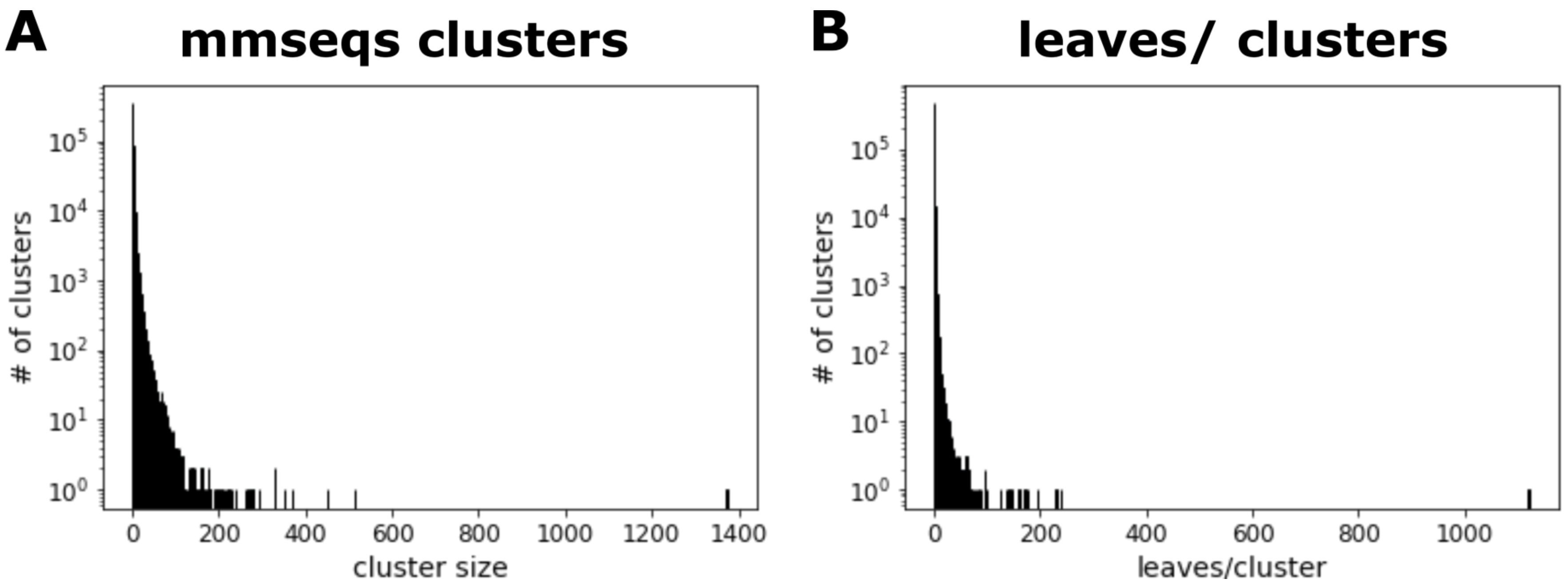

**Suppl. Fig.3: Profile annotation coverage across ICTV exemplar protein clusters**

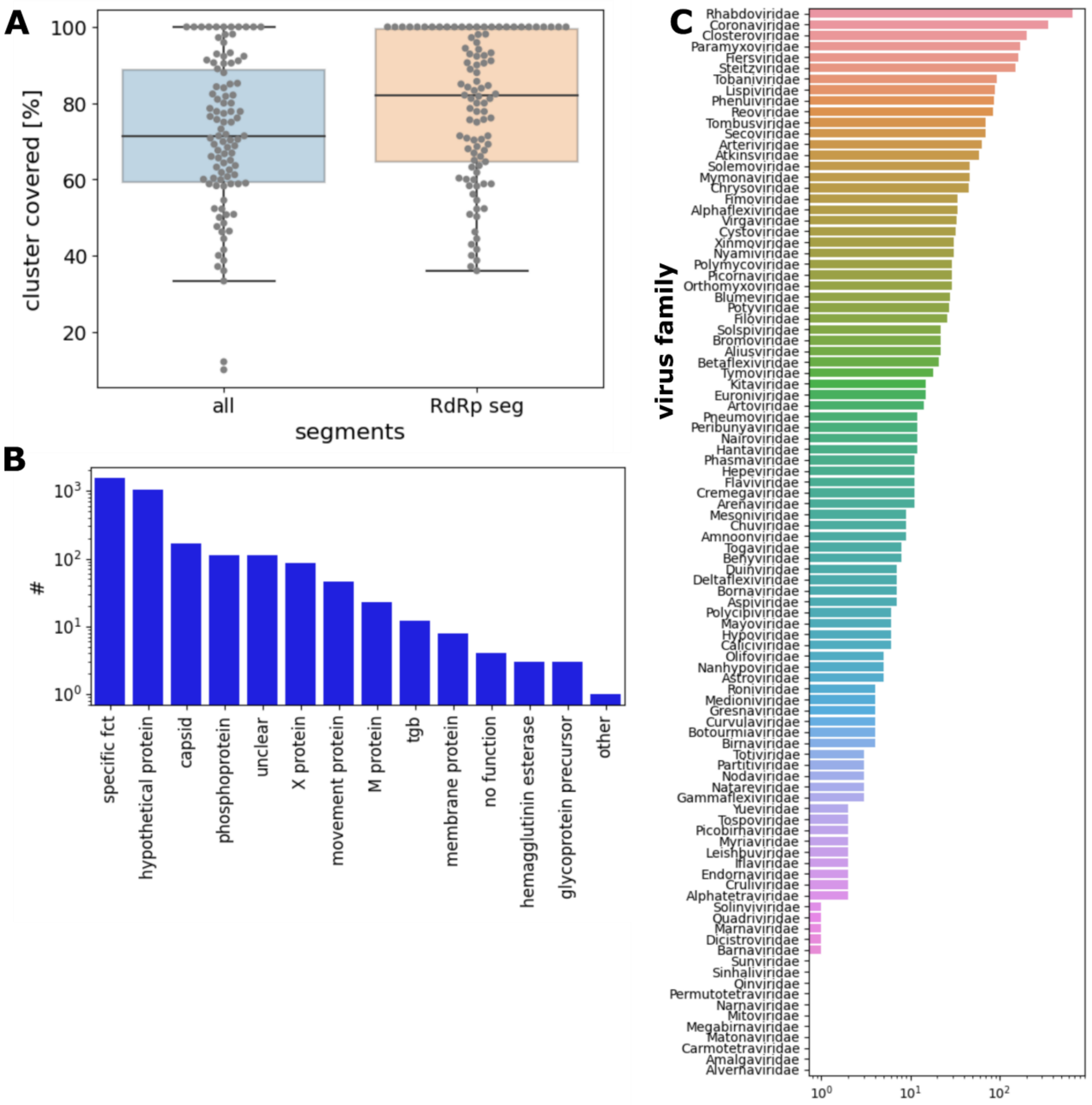

**Suppl.Fig.4: Characteristics of ORFan structures**

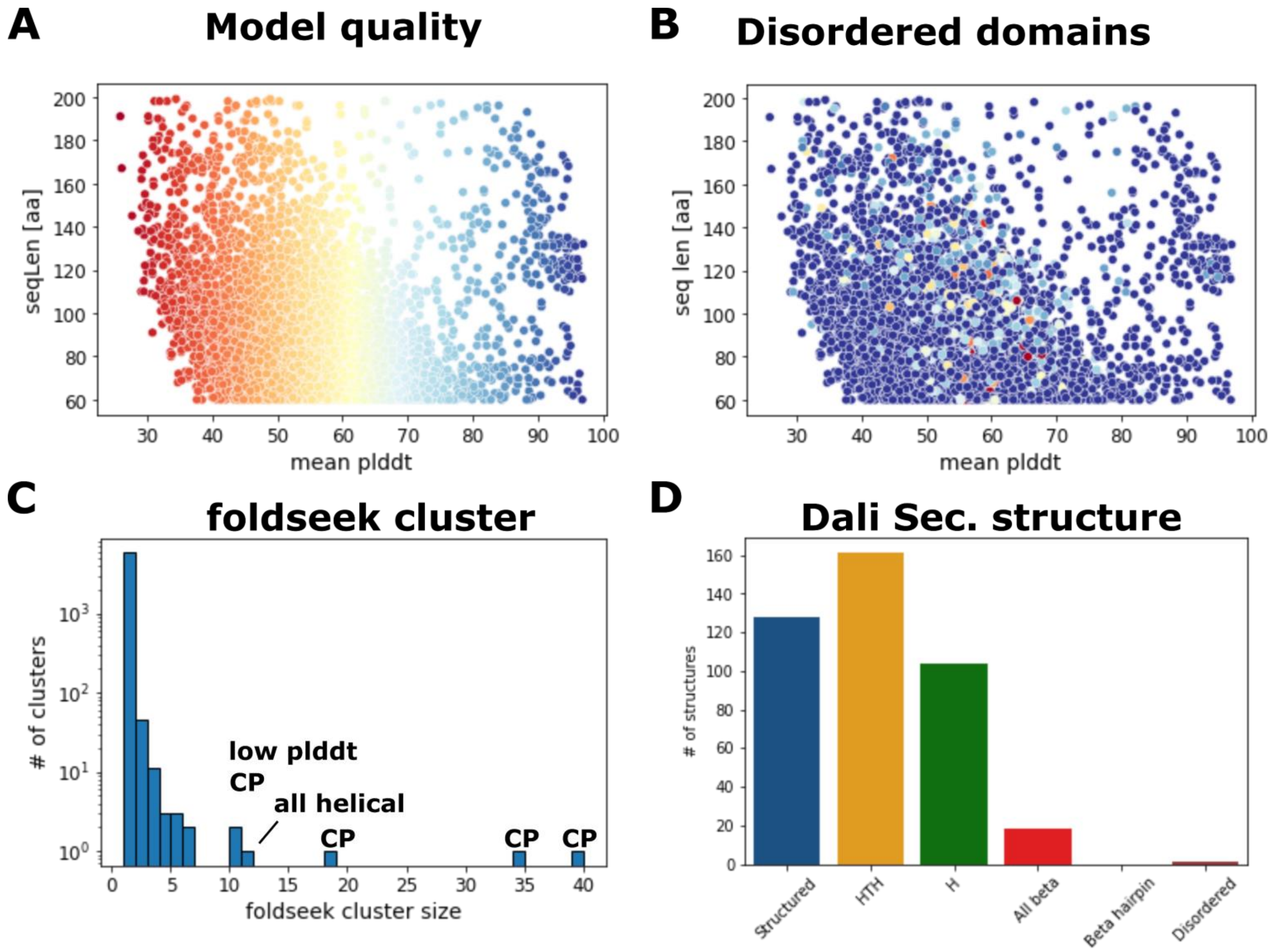

**Supp. Fig. 5: Characteristics of un-annotated domain structures**

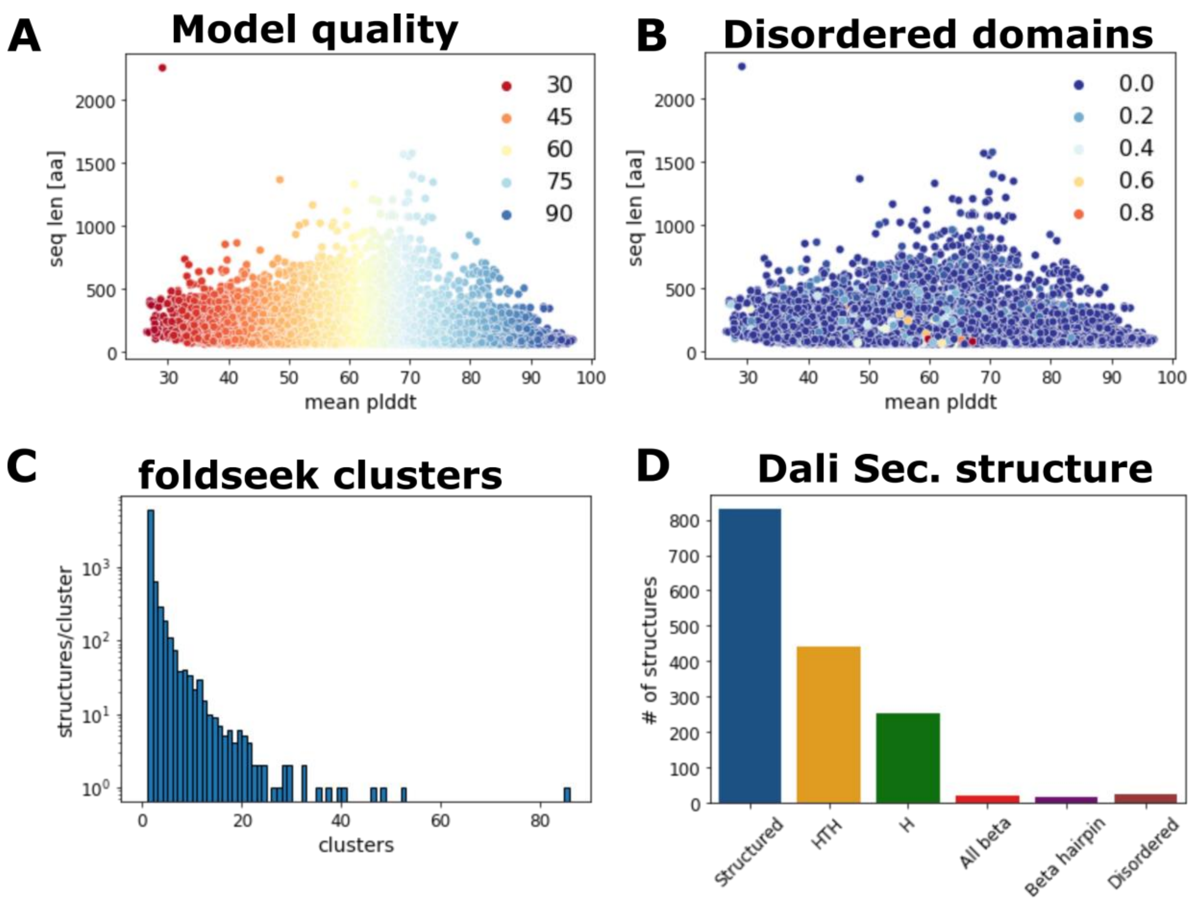

**Suppl. Fig. 6: Phylogenetic distribution of OOI and COI structure clusters**

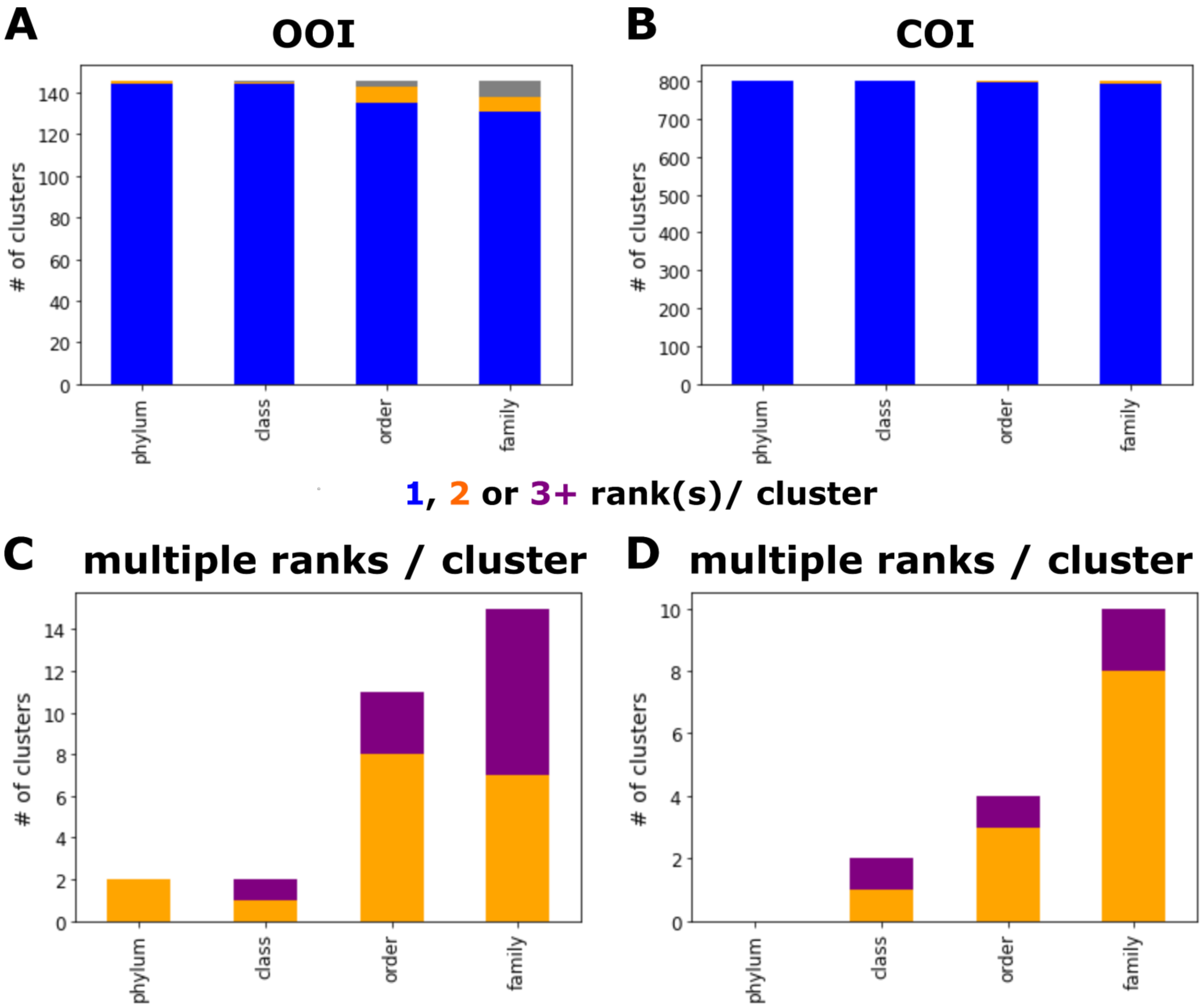

**Supp.Fig. 7: Structurome clustering**

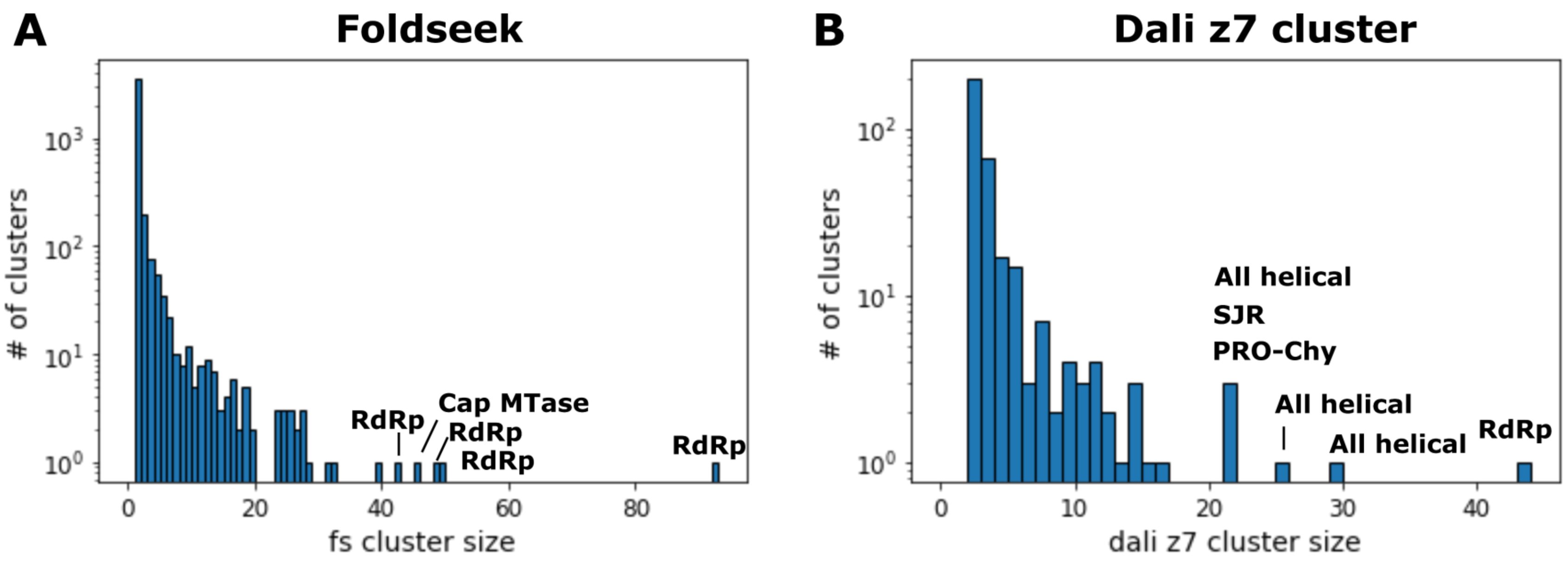

**Suppl. Fig 8: Structurome as structure-structure network**

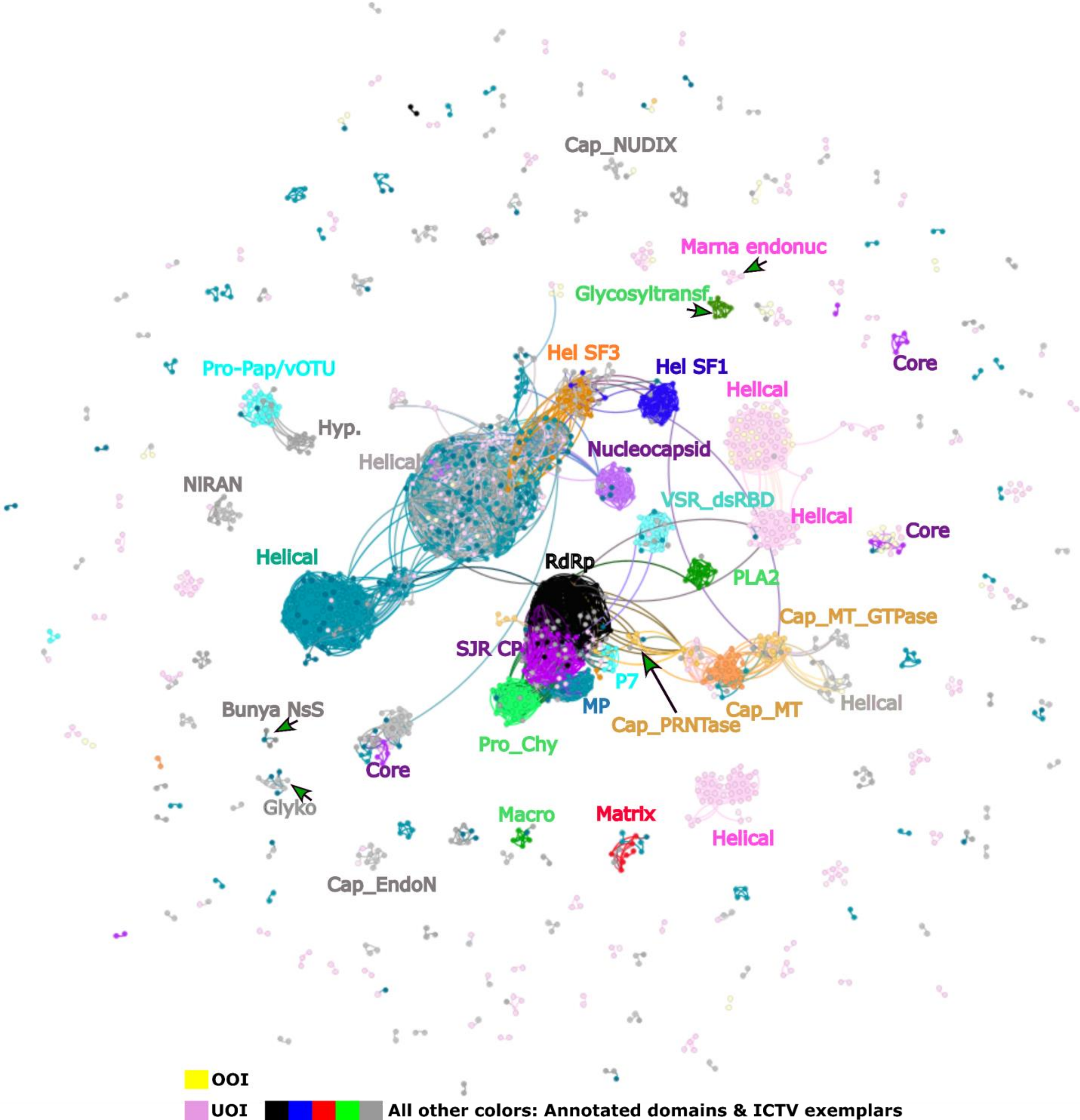

**Suppl. Fig. 9: Beta-barrel folds found in *f.0145***

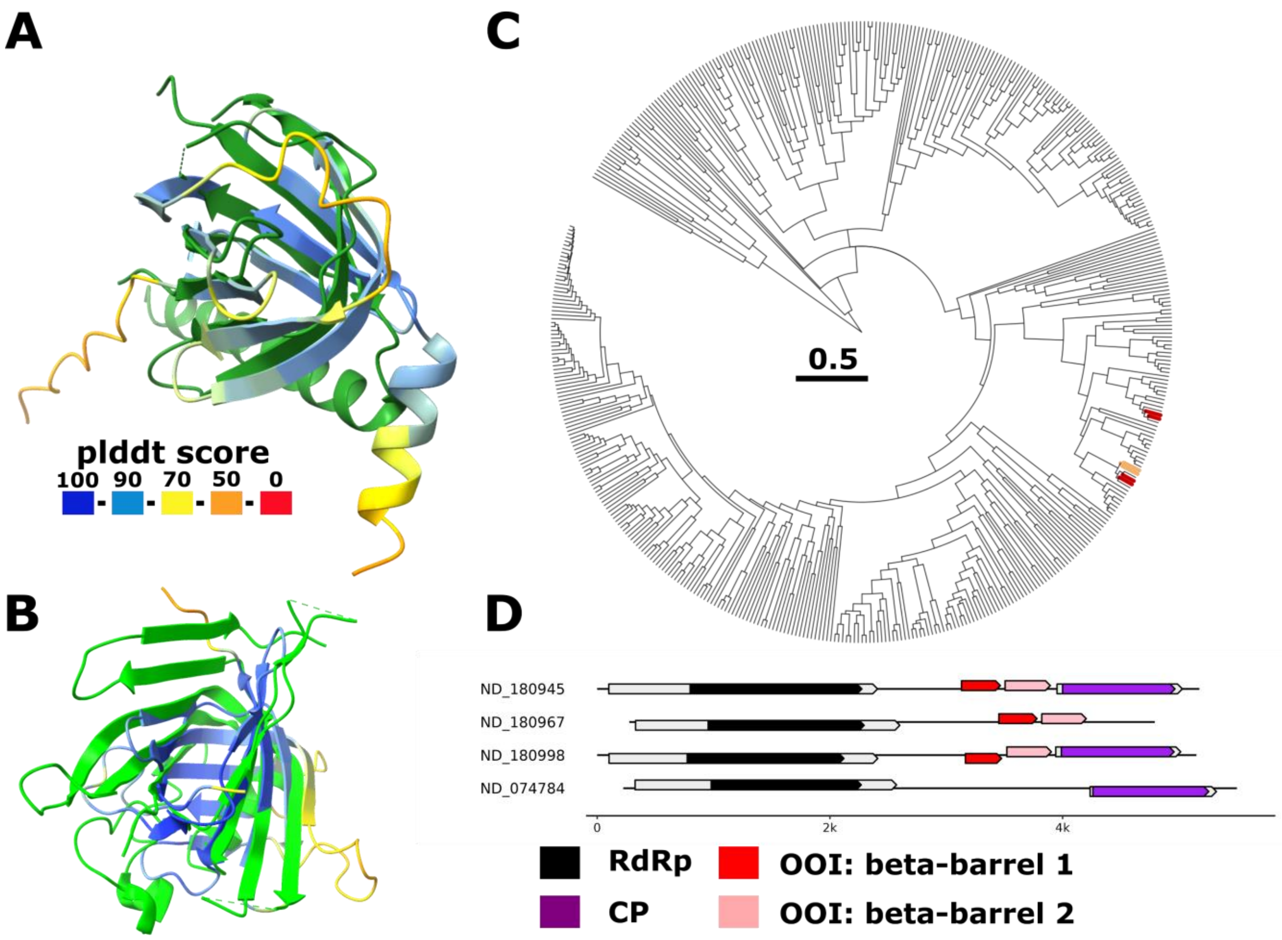

**Suppl. Fig. 10: Structure-guided alignment of viral core-p7 RBD-like folds and related cellular kinases**

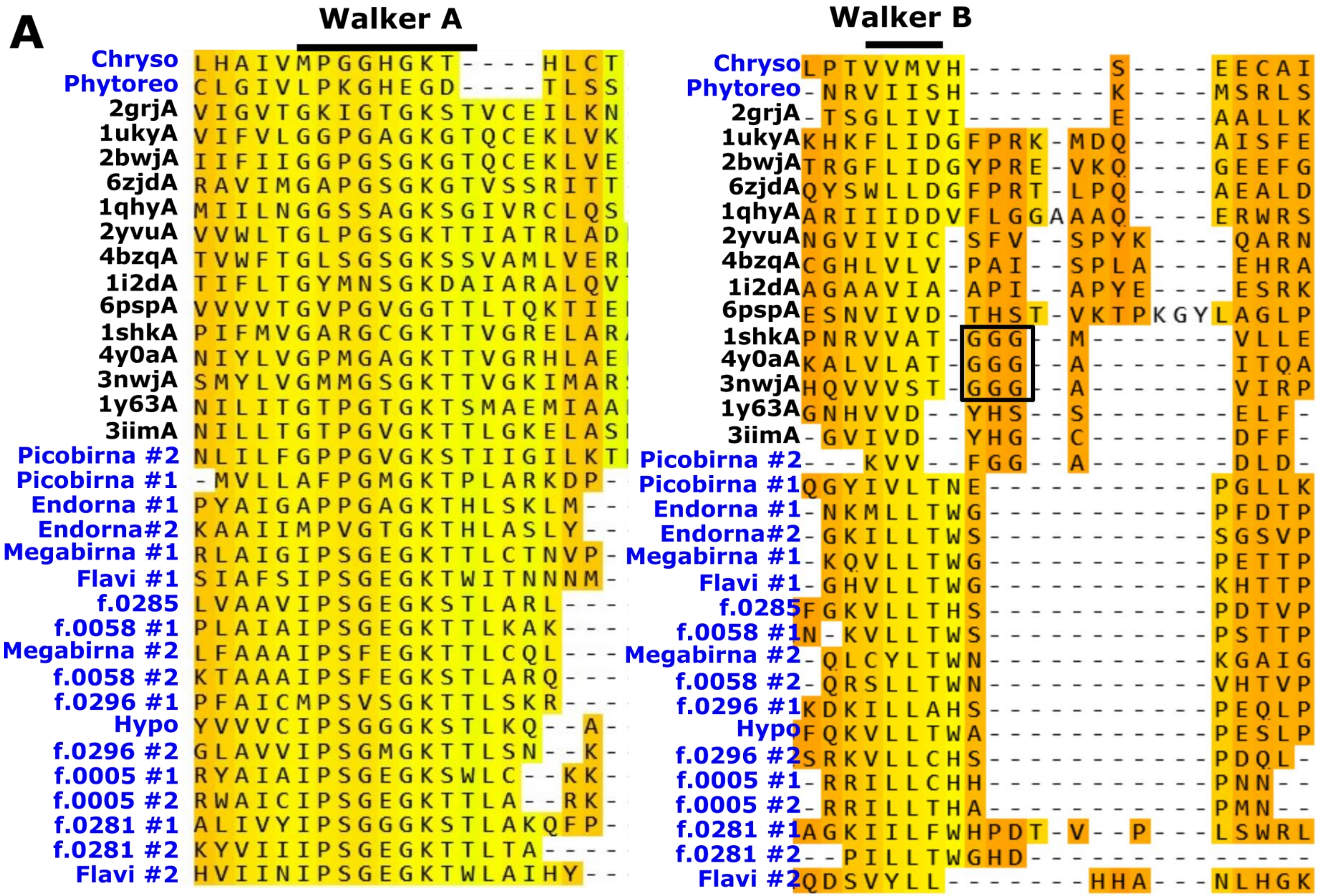

#### Suppl. Fig. 11: Viral core-p7 RBD-like folds from various viral families and genome organisation

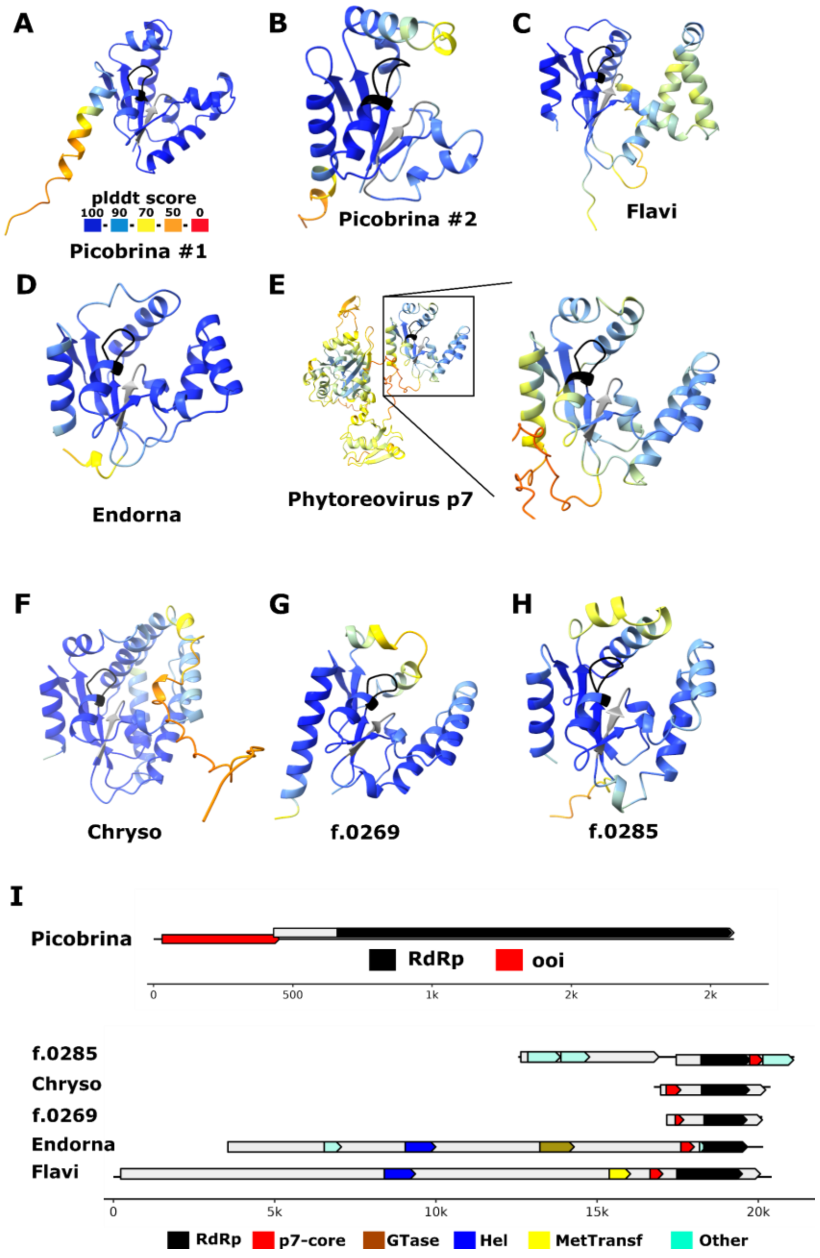

#### Suppl. Fig. 12: VSR-like RNA binding domain in f.0092.base-Permutotetra

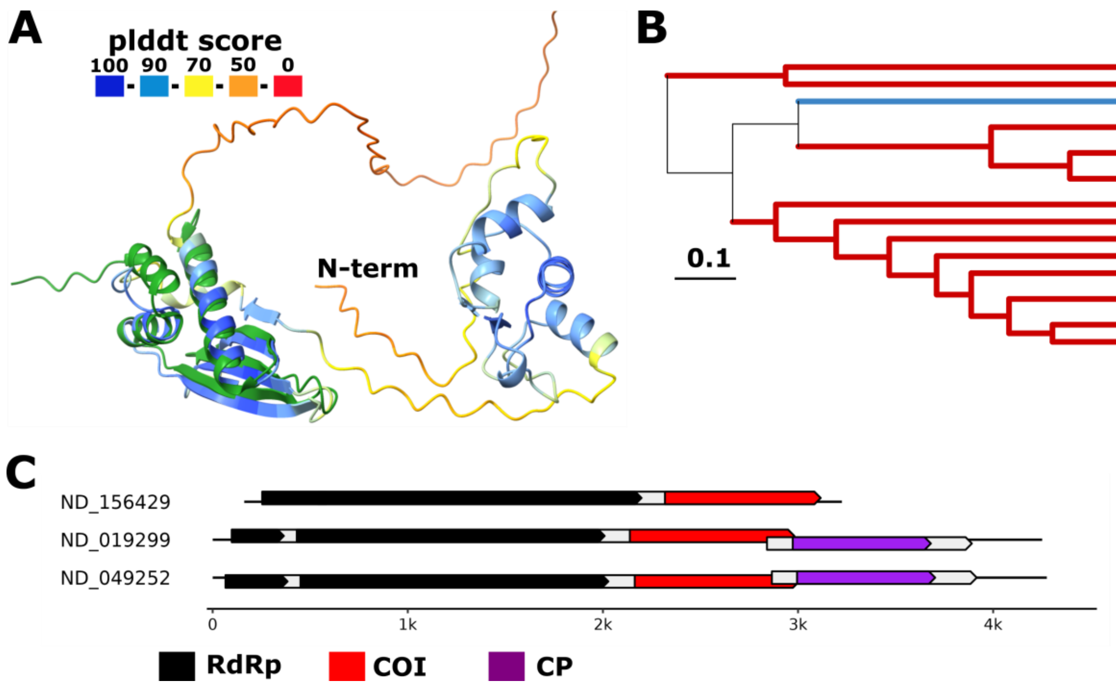

**Suppl. Fig. 13: Galactose binding domain in f.0008.base-Polycipi**

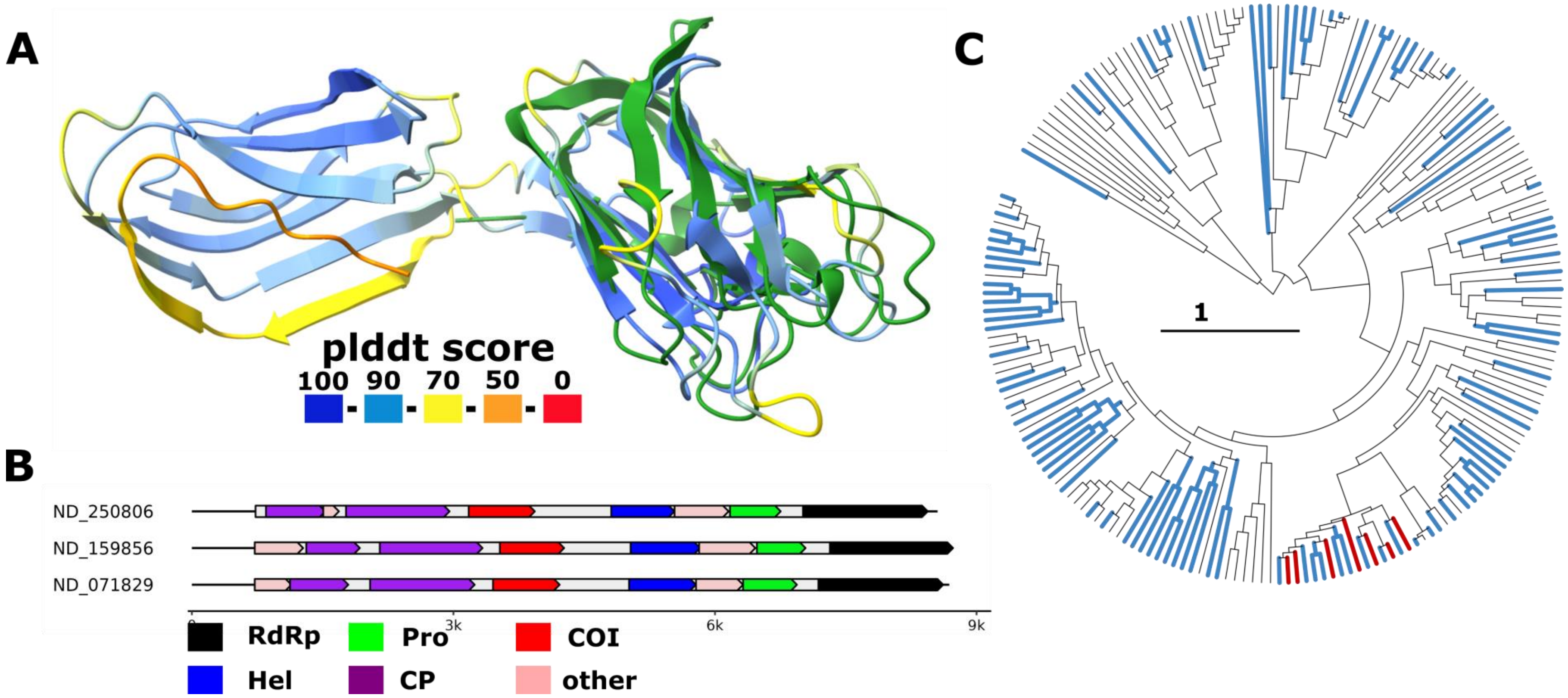

#### Suppl. Fig. 14: Exonuclease & methyltransferase in f.0181

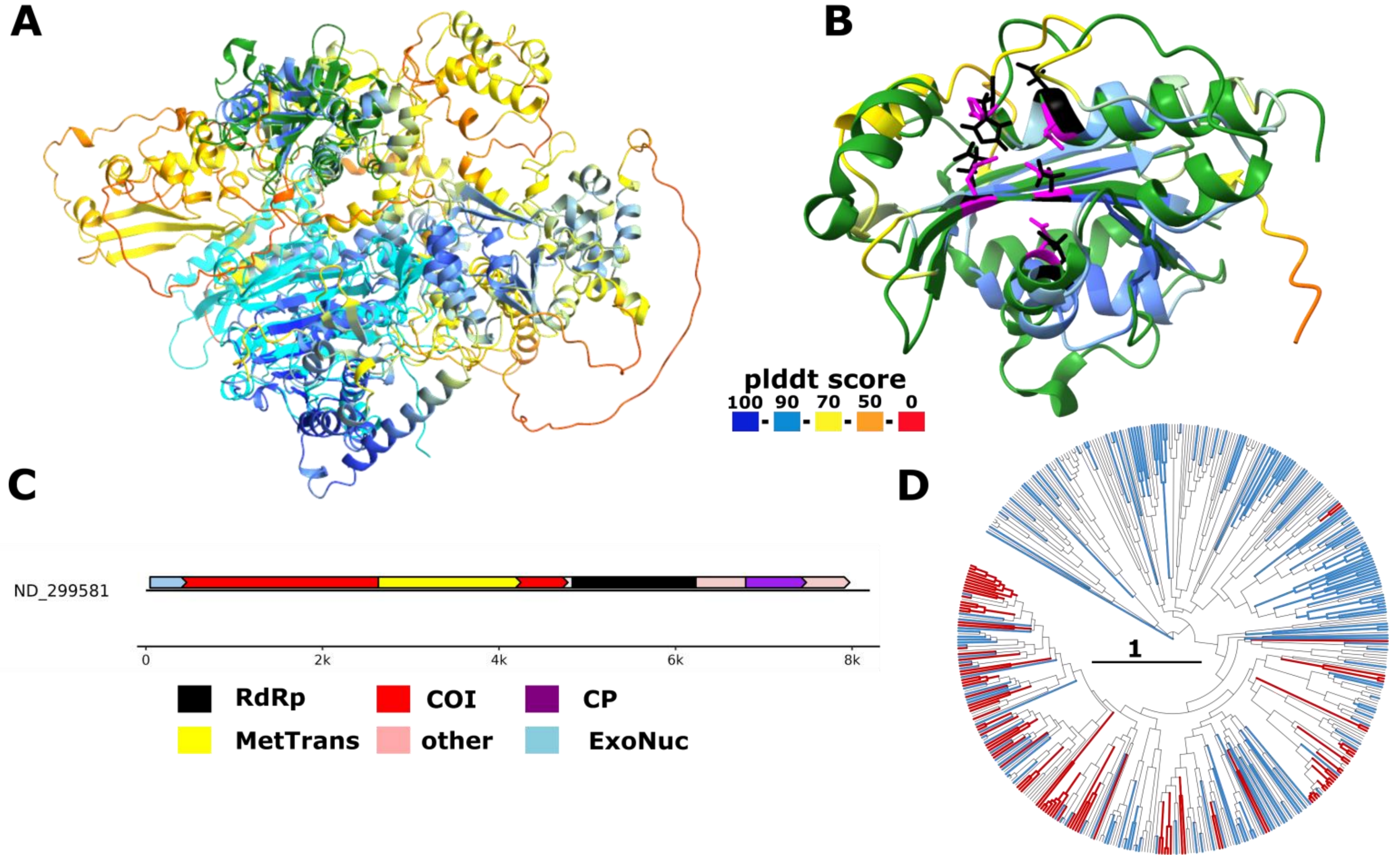

**Suppl. Fig. 15: Secovirus C-terminal domain of unknown function**

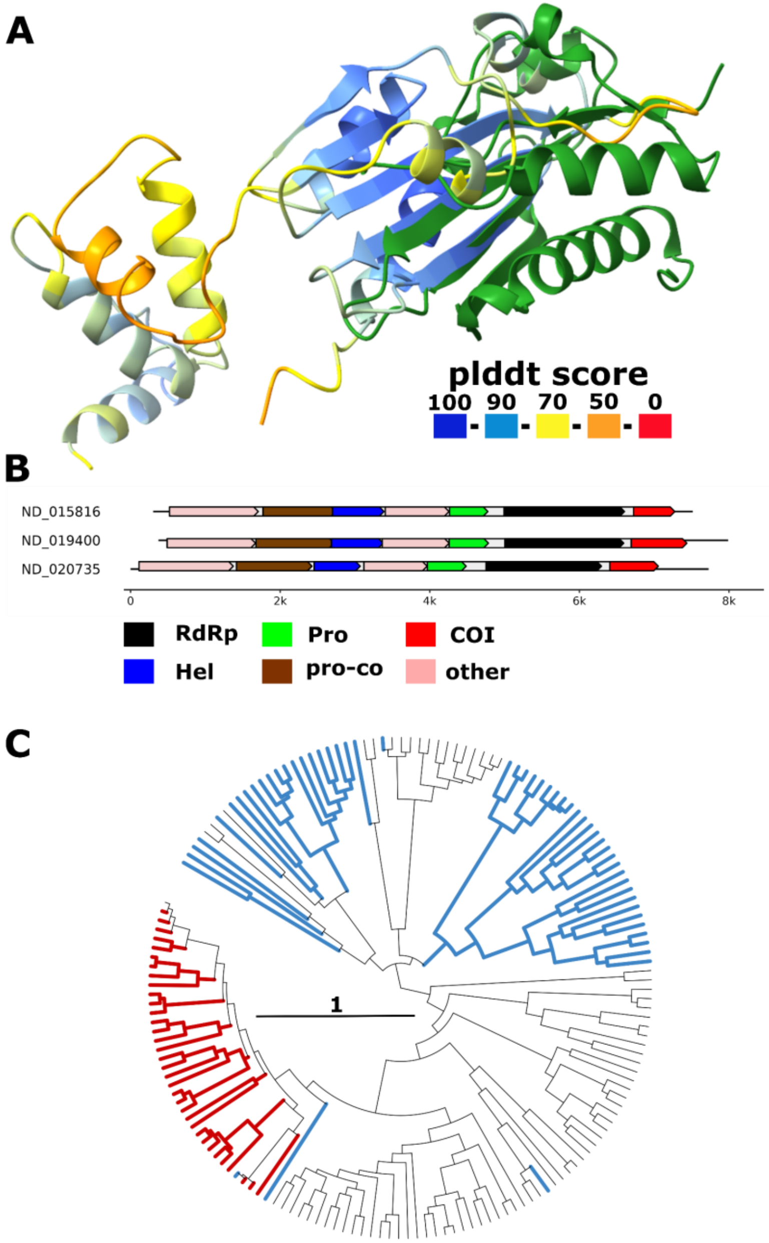

Suppl. Fig. 16: alpha-beta fold OOI found in *Rhabdoviridae*

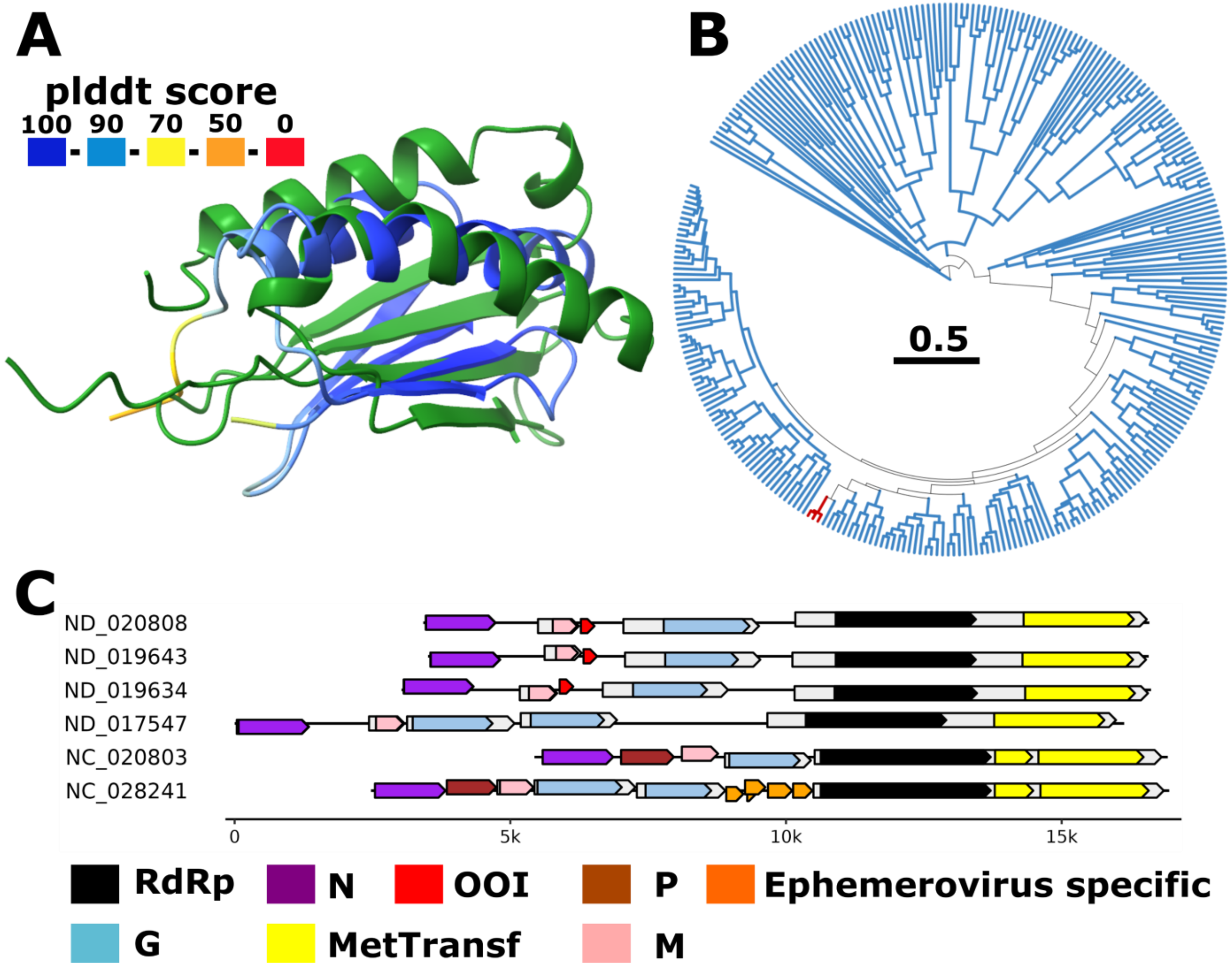

**Suppl. Fig. 17: vOTU in Deltaflexiviridae and related families**

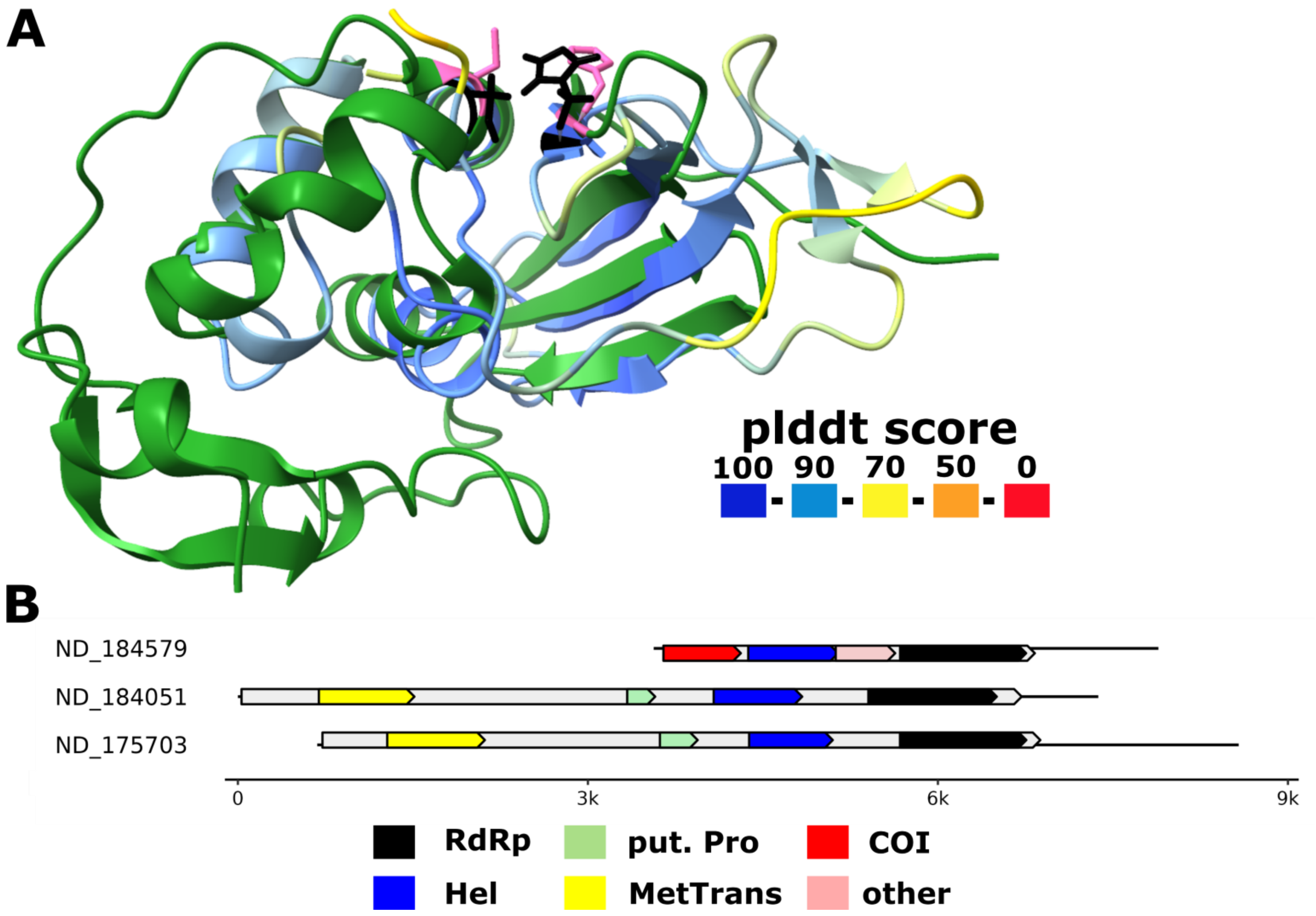

**Suppl. Fig. 18: Capsid protein in *f.0198***

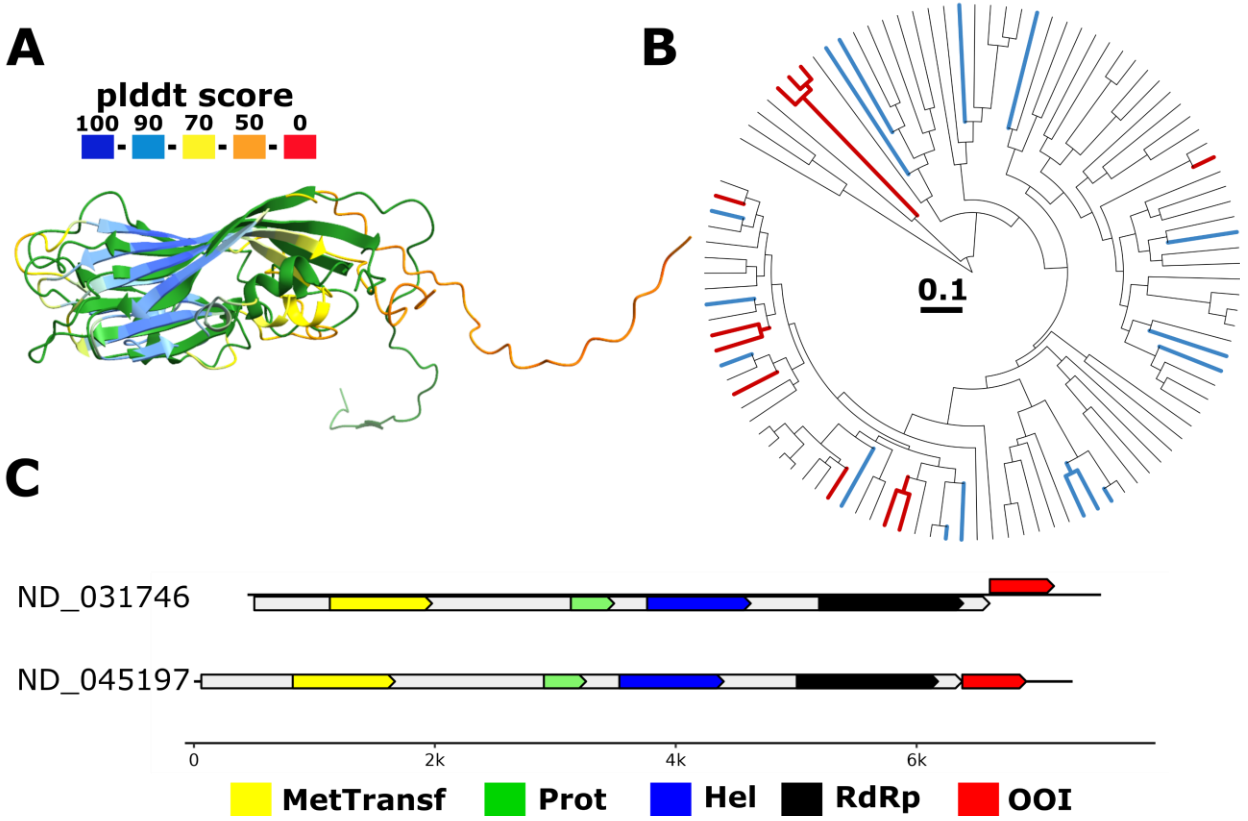

### Suppl. Fig. xx: Distinct domains per virus family

**A**

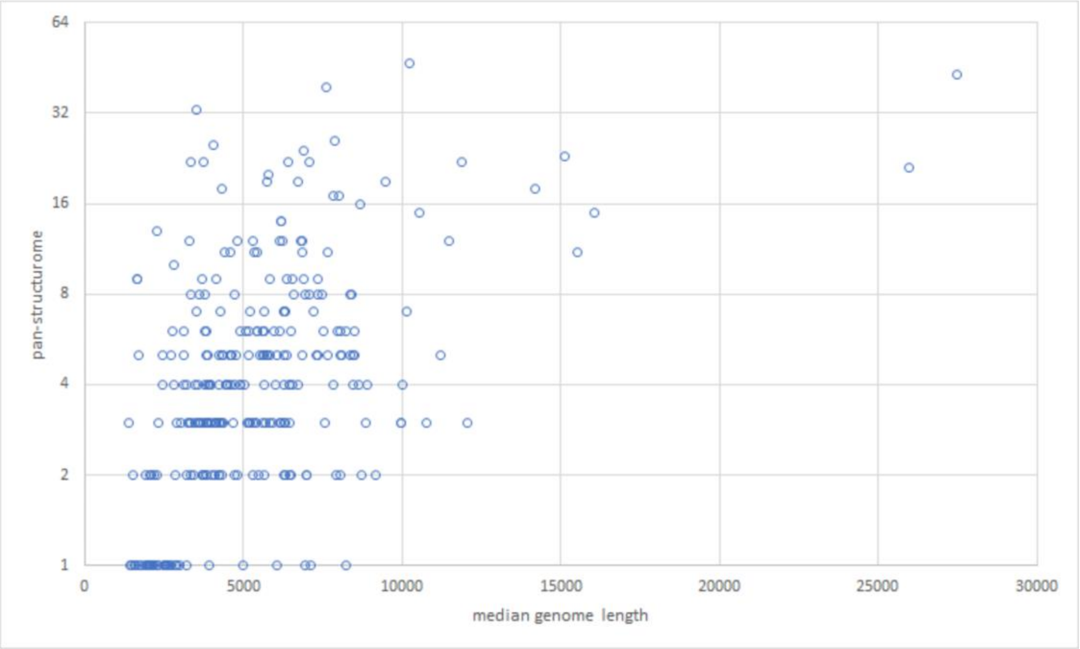


**B**

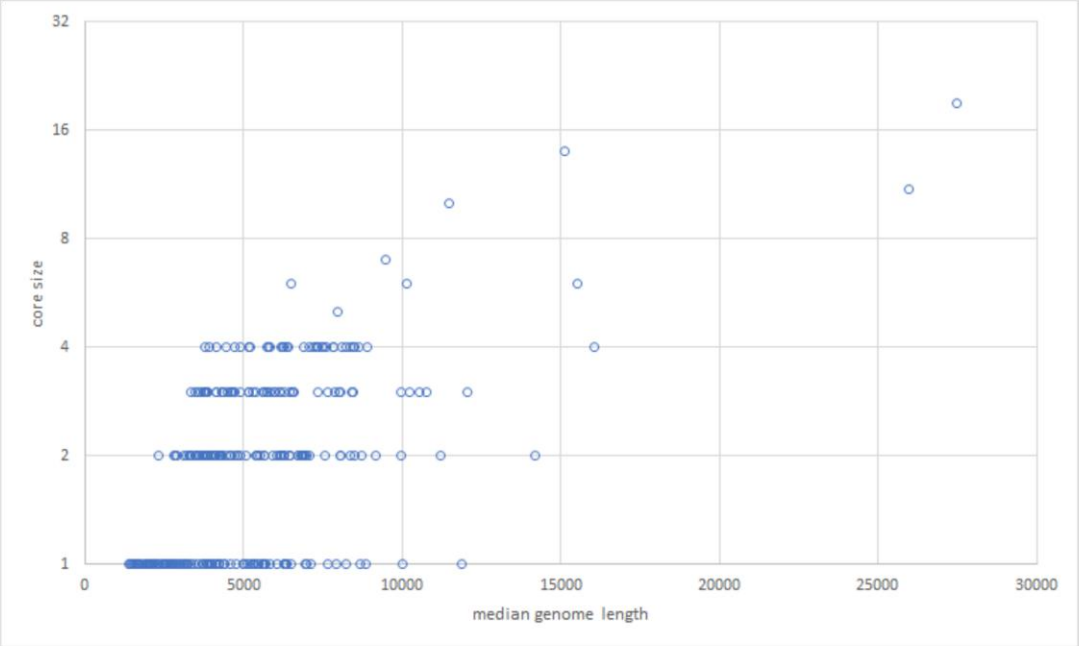
